# High resolution studies of DNA lesion bypass by human DNA polymerase δ holoenzymes

**DOI:** 10.1101/2022.06.30.498260

**Authors:** Rachel L. Dannenberg, Joseph A. Cardina, Kara G. Pytko, Mark Hedglin

## Abstract

During DNA replication, DNA lesions present in lagging strand templates are initially encountered by DNA polymerase δ (pol δ). The historical view for what transpires from these encounters is that replication of the afflicted lagging strand template abruptly stops, activating DNA damage tolerance (DDT) pathways that replicate the offending lesion and adjacent DNA sequence, allowing pol δ to resume downstream. However, qualitative studies observed that human pol δ is capable of replicating various DNA lesions, albeit to unknown extents, which raises issues regarding the roles of pol δ and DDT in the replication of DNA lesions. To address these issues, we re-constituted human lagging strand replication to quantitatively characterize initial encounters of pol δ holoenzymes with DNA lesions. The results indicate that pol δ holoenzymes support stable dNTP incorporation opposite and beyond multiple lesions and the extent of these activities depends on the lesion and pol δ proofreading. Furthermore, after encountering a given DNA lesion, subsequent dissociation of pol δ is distributed around the lesion and a portion of pol δ does not dissociate at all. The distributions of these events are dependent on the lesion and pol δ proofreading. These results challenge our understanding of DNA lesion replication and DDT.

## INTRODUCTION

In humans, like all eukaryotes, lagging strand DNA templates are primarily replicated by DNA polymerase δ (pol δ, **Figure 1**, *Top*), which is a member of the B-family of polymerases. Pol δ is comprised of four subunits; three accessory subunits (p50/POLD2, p66/POLD3, and p12/POLD4) and a catalytic subunit (p125/POLD1) that contains distinct active sites for DNA polymerase and 3′ → 5′ exonuclease (i.e., proofreading) activities. On its own, human pol δ is an inefficient and distributive DNA polymerase and must anchor to the processivity sliding clamp, proliferating cell nuclear antigen (PCNA), to form a pol δ holoenzyme with maximal efficiency and processivity (1). The highly conserved ring-shaped structure of PCNA has a central cavity large enough to encircle double-stranded DNA (dsDNA) and slide freely along it (2). Thus, association of pol δ with PCNA encircling a primer/template (P/T) junction effectively tethers the polymerase to DNA, substantially increasing the extent of continuous replication. The major single-strand DNA(ssDNA)-binding protein, replication protein A (RPA), engages the downstream template ssDNA that is to be replicated, preventing its degradation by cellular nucleases and formation of secondary DNA substrates that are prohibitive to DNA replication (3). Furthermore, upon dissociation of pol δ from a P/T junction, RPA prevents diffusion of PCNA along the adjacent 5′ ssDNA overhang (4,5).

**Figure 1.**
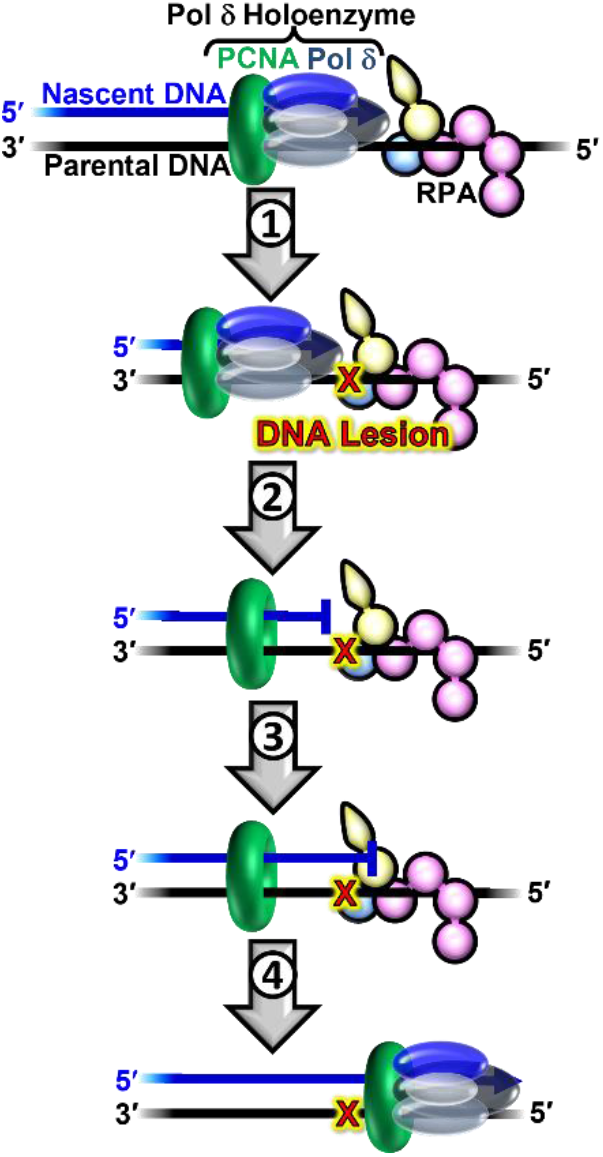
DNA damage tolerance in lagging strand templates. At the top, a progressing pol δ holoenzyme (pol δ + PCNA) is depicted replicating a lagging strand template engaged by RPA. **1**) A progressing pol δ holoenzyme encounters a DNA lesion in a lagging strand template. **2**) Pol d rapidly and passively dissociates into solution, leaving PCNA and RPA behind on the DNA. Pol d may reiteratively associate and dissociate to/from the resident PCNA encircling the stalled P/T junction cannot support stable dNTP incorporation opposite the offending DNA lesion. **3**) The stalled P/T junction activates one or more DNA damage tolerance pathway(s) that are ultimately responsible for the insertion of a dNTP opposite the lesion (insertion), extension of the nascent DNA 1 nt downstream of the lesion (extension), and possibly further extension of the nascent DNA >1 nt downstream of the lesion (elongation). **4**) After DDT is complete, replication by pol δ holoenzymes may resume downstream of the lesion. In this view, only DDT is responsible for the replication of a DNA lesion, and, hence, pol δ does not contribute to the fidelity of replicating DNA lesions.

As the primary lagging strand DNA polymerase, pol δ is the first to encounter lagging strand template nucleotides that have been damaged by covalent modifications. These damaging modifications, often referred to as DNA lesions, arise from exposure of genomic DNA to reactive metabolites and environmental mutagens. Given the highly stringent DNA polymerase activity of human pol δ along with its robust, intrinsic proofreading activity, the historical view (**Figure 1**) for what transpires upon human pol δ encountering a DNA lesion is that pol δ dissociates into solution, leaving PCNA and RPA behind at the aborted P/T junction. Pol δ may re-iteratively associate and dissociate from the resident PCNA, but it cannot support stable insertion of a dNTP opposite the lesion. Consequently, replication of the lagging strand template stalls, activating DNA damage tolerance (DDT) pathways that are ultimately responsible for insertion of a dNTP opposite the lesion (i.e., insertion), extension of the nascent DNA 1 nucleotide (nt) downstream of the lesion (i.e., extension), and possibly further elongation of the nascent DNA > 1 nt downstream of the lesion (i.e., elongation). After DDT, replication by pol δ holoenzymes may resume downstream of the lesion (6,7). However, over the last 15 years or so, numerous qualitative studies from independent groups (8-14) observed that human pol δ is capable of replicating various DNA lesions, albeit to unknown extents, which raises an issue of whether pol δ is directly involved in DDT and hence a major player in the fidelity of replicating DNA lesions. This issue has critical implications for our understanding of when, how, and why DDT is activated. To address this issue, we re-constituted human lagging strand replication at physiological pH, ionic strength, and dNTP concentrations to quantitatively characterize, at single nucleotide resolution, the initial encounters of pol δ holoenzymes with downstream DNA lesions. In short, a DNA lesion ≥ 9 nt downstream of a P/T junction is encountered only once and only by a progressing pol δ holoenzyme, rather than pol δ alone. To the best of our knowledge, comparable studies on human lagging strand replication have yet to be reported. The results indicate that human pol δ holoenzymes support stable dNTP incorporation opposite and beyond multiple lesions and the extent of these activities depends on the identity of the lesion and the ability to proofread intrinsically (as opposed to extrinsically). Furthermore, the results indicate that, after encountering a given DNA lesion, subsequent dissociation of pol δ does not occur at a uniform site relative to the lesion. Rather, pol δ dissociation events are distributed around the lesion and a portion of pol δ does not dissociate at all. The distributions of these events are dependent on the identity of the lesion and the ability to proofread intrinsically. These results challenge our understanding of DNA lesion replication and its fidelity as well as the activation and function of DDT.

## Materials and Methods

### Recombinant Human Proteins

Human RPA, Cy5-PCNA, RFC, and pol δ (exonuclease-deficient and wild-type) were obtained as previously described (15,16). The concentration of active RPA was determined via a FRET-based activity assay as described previously (17).

### Oligonucleotides

Oligonucleotides were synthesized by Integrated DNA Technologies (Coralville, IA) or Bio-Synthesis (Lewisville, TX) and purified on denaturing polyacrylamide gels. The concentrations of unlabeled DNAs were determined from the absorbance at 260 nm using the calculated extinction coefficients. The concentrations of Cy5-labeled DNAs were determined from the extinction coefficient at 650 nm for Cy5 (ε_650_ = 250,000 M^−1^cm^−1^). The concentrations of Cy3-labeled DNAs were determined from the extinction coefficient at 550 nm for Cy3 (ε_650_ = 125,000 M^−1^cm^−1^). For annealing two single strand DNAs, the primer and corresponding complementary template strands were mixed in equimolar amounts in 1X Annealing Buffer (10 mM TrisHCl, pH 8.0, 100 mM NaCl, 1 mM EDTA), heated to 95 °C for 5 minutes, and allowed to slowly cool to room temperature.

### Primer Extension Assays

All primer extension experiments were performed at 25 °C in an assay buffer consisting of 1X Replication Buffer supplemented with 1 mM DTT and 1 mM ATP. For all experiments, the final ionic strength was adjusted to 230 mM by addition of appropriate amounts of KOAc and samples are protected from light whenever possible. All reagents, substrate, and protein concentrations listed are final reaction concentrations. First, 250 nM Cy5-labeled P/T DNA (**Figure S1**) is preincubated with 1 μM Neutravidin. Next, RPA (750 nM heterotrimer) is added and the resultant mixture is allowed to equilibrate for 5 min. PCNA (250 nM homotrimer), ATP (1 mM), and RFC (250 nM heteropentamer) are then added in succession and the resultant mixture is incubated for 5 min. Finally, dNTPs (46 μM dATP, 9.7 μM dGTP, 48 μM dCTP, 67 μM dTTP) are added and DNA synthesis is initiated by the addition of limiting pol δ (either 8.8 nM wild-type or 35 nM exonuclease-deficient heterotetramer). The concentration of each dNTP utilized is within the physiological range observed in dividing human cells (24 ± 22 μM dATP, 5.2 ± 4.5 μM dGTP, 29 ± 19 μM dCTP, 37 ± 30 μM dTTP) (18). At variable times, aliquots of the reaction were removed, quenched with 62.5 mM EDTA, pH 8.0, 2 M Urea, 50% formamide supplemented with 0.01% (wt/vol) tracking dyes. Primer extension products were resolved on 16% sequencing gels. Before loading onto gel, quenched samples were heated at 95 °C for 5 min and immediately chilled in ice water for 5 min. Gel images were obtained on a Typhoon Model 9410 imager. The fluorescence intensity in each band on a gel was quantified with ImageQuant (GE Healthcare) and the fluorescence intensity of each DNA band within a given lane was converted to concentration by first dividing its intensity by the sum of the intensities for all of the species present in the respective lane and then multiplying the resultant fraction by the concentration of P/T DNA (250 nM). Within a given lane, the probability of incorporation, *P*_i_, for each dNTP incorporation step, *i*, after *i*_1_ was calculated as described previously (1,19). The insertion probability is the probability of dNTP incorporation opposite a DNA lesion or the corresponding native nucleotide at dNTP incorporation step *i* and is equal to *P*_i_. The insertion efficiency was calculated by dividing the insertion probability for a given DNA lesion by the insertion probability for the corresponding native nucleotide in the same sequence context and then multiplying the resultant quotient by 100%. The extension probability is the probability of dNTP incorporation 1 nt downstream of a DNA lesion or the corresponding native nucleotide at dNTP incorporation step *i* and is equal to *P*_i + 1_. The extension efficiency was calculated by dividing the extension probability for a given DNA lesion by the extension probability for the corresponding native nucleotide in the same sequence context and then multiplying the resultant quotient by 100%. The bypass probability for a given DNA lesion or the corresponding native nucleotide at dNTP insertion step, *i*, was calculated by multiplying *P*_i_ by *P*_i+1._ The bypass probability represents that probability of dNTP incorporation opposite a DNA lesion or the corresponding native nucleotide at dNTP incorporation step *i*, and the next dNTP incorporation step downstream (*i* +1). The bypass efficiency was calculated by dividing the bypass probability for a given DNA lesion by the bypass probability for the corresponding native nucleotide in the same sequence context and then multiplying the resultant quotient by 100%. Upon encountering a given DNA lesion or the corresponding native nucleotide at dNTP incorporation step, *i*, the fraction of pol δ that subsequently dissociates at dNTP incorporation step *i* or any dNTP incorporation step downstream is defined as the band intensity at that dNTP incorporation step divided by the sum of the band intensity at that dNTP incorporation step and the band intensities for all longer primer extension products. These values are utilized to determine the distribution (%) of pol δ dissociation events that occur after the initial encounter with a DNA lesion or the corresponding native nucleotide at dNTP incorporation step *i*. Only data points that are less than 20% of the reaction progress (based on the accumulation of primer extension products) were plotted as a function of time and analyzed with Kaleidagraph (Synergy). Within the linear phase of primer extension, the *P*_i_ values, the variables calculated from *P*_i_ values, and all parameters discussed above remained constant with incubation time. For a given dNTP incorporation step, the *P*_i_ values within the linear phase of primer extension, the variables calculated from these *P*_i_ values, and all parameters discussed above were each fit to a flat line where the y-intercept reflects the average value.

## RESULTS

### Strategy to monitor progression of human pol δ holoenzymes

The approach utilizes P/T DNA substrates (**Figure S1**) that mimic nascent P/T junctions on a lagging strand. Each P/T DNA is comprised of a 62-mer template strand annealed to a 29-mer primer strand that contains a biotin at the 5′ terminus and an internal Cy5 dye label 4 nt from the 5′ terminus. When pre-bound to Neutravidin, the biotin prevents loaded PCNA from sliding off the dsDNA end of the substrate. The lengths (29 base pairs, bp) of the dsDNA regions are identical and in agreement with the requirements for assembly of a single PCNA ring onto DNA by RFC (4,5,16). The lengths (33 nt) of the ssDNA regions adjacent to the 3′ end of the P/T junctions are identical and accommodate 1 RPA molecule (20-22). P/T DNA is pre-saturated with Neutravidin and RPA and then PCNA is assembled onto all P/T junctions by RFC and stabilized by RPA and Neutravidin/biotin blocks that prohibit PCNA from diffusing off the P/T DNA (**Figure S2**) (4,5). Finally, primer extension (i.e., dNTP incorporation) is initiated by the addition of limiting pol δ (**Figure 2A**) and Cy5-labeled DNA products are resolved on a denaturing polyacrylamide gel (**Figure 2B**), visualized on a fluorescent imager, and quantified.

**Figure 2.**
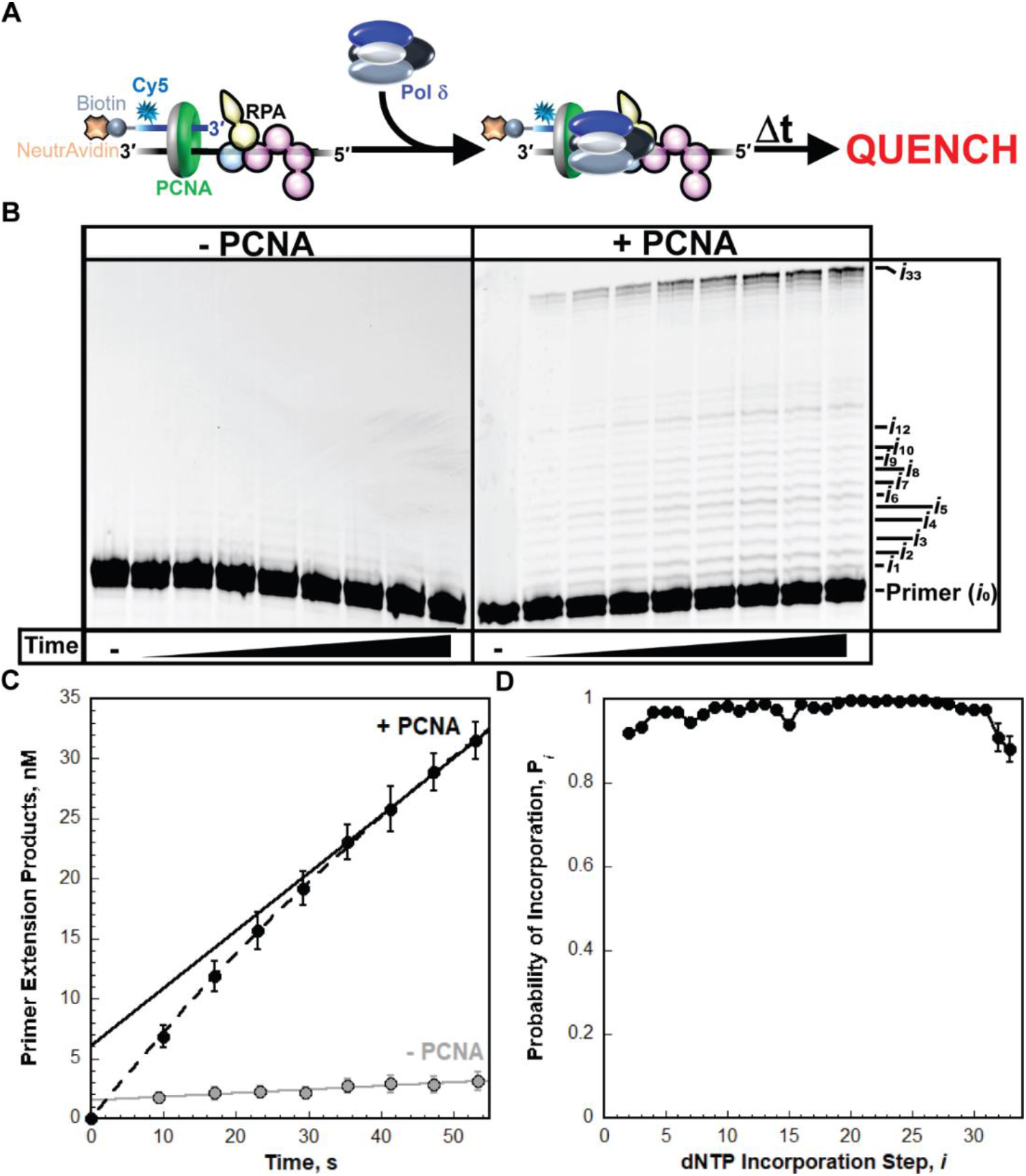
Replication by pol δ holoenzymes. (**A**) Schematic representation of the experiment performed to monitor primer extension by pol δ holoenzymes during a single binding encounter with a P/T DNA substrate. PCNA is assembled onto a native (i.e., undamaged) DNA substrate (BioCy5P/T, **Figure S1**) by the successive addition of RPA, PCNA, ATP, and RFC. Physiological concentrations of dNTPs are then added, and synthesis is initiated by the addition of limiting pol δ. (**B**) 16% denaturing sequencing gel of the primer extension products. The incorporation step (*i*) for certain primer extension products (*i*_*1*_ to *i*_10_, *i*_12_, and *i*_33_) is indicated on the far right. Shown on the left and the right are representative gels of primer extension by pol δ observed in the absence (“-PCNA”) and presence (“+ PCNA”) of PCNA, respectively. (**C**) Quantification of the (total) primer extension products. Each data point represents the average ± S.E.M. of 3 independent experiments. Data is plotted as a function of time (after the addition of pol δ) and display “burst” kinetics. Data points within the “linear” phase are fit to a linear regression where the Y-intercept (in nM) represents the amplitude of the “burst” phase, and the slope represents the initial velocity (in nM/min) of the linear phase. Data for experiments carried out in the absence (“-PCNA”) and presence (“+PCNA”) of PCNA are displayed in grey and black, respectively. Data for experiments carried out in the presence of PCNA are fit to a burst + linear phase kinetic model (dashed line) only for visualizing the conformity of the linear phases for each fit. (**D**) Processivity of pol δ holoenzymes. The probability of incorporation (*P*_i_) for each dNTP incorporation step (*i*) beyond the first incorporation step is calculated as described in *Experimental Procedures. P*_i_ values observed in the presence of PCNA are plotted as a function of the dNTP incorporation step, *i*, and each data point represents the average ± S.E.M. of 3 independent experiments. Data is fit to an interpolation only for observation. Error bars are present for all data points on all plots but may be smaller than the data point.

Under the conditions of the assay (physiological pH, ionic strength, and concentration of each dNTP), primer extension on a control (i.e., native/undamaged) P/T DNA (BioCy5P/T, **Figure S1**) is severely limited in the absence of PCNA and not observed beyond the 5^th^ dNTP incorporation step (i.e., *i*_5_) (**Figure 2B**, “-PCNA”) whereas significant primer extension is observed in the presence of PCNA up to and including the last dNTP incorporation step (*i*_33_) (**Figure 2B**, “+PCNA”). In the absence of PCNA, only 1.249 ± 0.3149 % of the primer is extended over the time period monitored (60 s) compared to 12.60 ± 0.6132 % in the presence of PCNA (**Figure 2C**). These observations agree with the inability of pol δ alone to form a stable complex with native P/T DNA (1,23). Altogether, the results from **Figure 2A – 2C** indicate that nearly all DNA synthesis (> 90%) observed in the presence of PCNA is carried out by pol δ holoenzymes and only pol δ holoenzymes are responsible for primer extension beyond the 5^th^ dNTP incorporation step (i.e., *i* ≥ 5).

All primer extension assays reported in this study were performed in the presence of a large excess of P/T DNA over pol δ and only monitor ≤ 20% of the reaction such that once a primer is extended and the associated pol δ subsequently disengages, the probability that the extended primer will be utilized again is negligible. Rather, the dissociated pol δ engages another, previously unused primer. In other words, the observed primer extension products reflect a single cycle (i.e., single pass, single hit, etc.) of DNA synthesis. Appropriate single hit conditions are operating for any pol δ:P/T DNA ratios when the probabilities of dNTP incorporation (*P*_i_) remain constant with incubation time, as depicted in **Figure S3** for the BioCy5P/T DNA substrate in the presence of PCNA (1,19,24,25). This condition was met for all *P*_i_ values reported in this study. For a given dNTP incorporation step, *i*, the probability of dNTP incorporation, *P*_i_, represents the likelihood that pol δ will incorporate a dNTP rather than dissociate. For the BioCy5P/T DNA substrate, the *P*_i_ values observed in the presence of PCNA (**Figure 2D**) are high and range from 0.998 ± 0.00118 for the 23^rd^ dNTP incorporation (*i*_23_) to 0.881 ± 0.0308 for the last dNTP incorporation step (*i* = 33). Maximal *P*_i_ values (≥ 0.990) are observed beginning at *i*_16_ and are maintained until *i*_27_, after which *P*_i_ drops off, particularly at *i*_32_ and *i*_33_, as progressing pol δ holoenzymes dissociate due to the severely diminished length (2 nt) of the single strand template. Importantly, the distribution and range of observed *P*_i_ values are in excellent agreement with values reported in previous studies on the same P/T DNA substrate where only DNA synthesis by pol δ holoenzymes is observed (1,19). This re-affirms that > 90%, if not all, DNA synthesis observed in the presence of PCNA was carried out by pol δ holoenzymes, as opposed to pol δ alone. All *P*_i_ values observed for the BioCy5P/T DNA substrate in the presence of PCNA are less than 1.0 (**Figure 2D**) indicating that a proportion of progressing pol δ holoenzymes dissociate at each successive dNTP incorporation step. This behavior of pol δ holoenzymes on the native P/T DNA agrees very well with the extensively documented behavior of human pol δ holoenzymes (1,19,26-32). This assay was utilized in the present study to directly compare the progression of human pol δ holoenzymes on the native/undamaged BioCy5P/T DNA substrate (**Figure S1**) to that observed on damaged P/T DNA substrates that are identical to the BioCy5P/T DNA except that a single nt ≥ 9 nt downstream of the P/T junction is altered by a chemical modification(s), i.e., DNA lesion(s) (**Figure S1**). The DNA lesions examined in the present study are prominent in cells exposed to oxidizing or alkylating agents. In this “running start” setup, dNTP incorporation initiates upstream of a DNA lesion and, hence, the DNA lesion is encountered by progressing pol δ holoenzymes that have a “running start.” Stable assembly (i.e., loading) of PCNA onto a P/T junction was not affected by any of the DNA lesions examined in the present study (**Figure S2**). Thus, any observed effect on the DNA synthesis activity of assembled pol δ holoenzymes is not attributable to the amount of PCNA loaded onto a P/T DNA junction with a DNA lesion ≥ 9 nt downstream.

### The effect of oxidative DNA lesions on the progression of human pol δ holoenzymes

First, we examined the effect of 7,8-dihydro-8-oxoguanine (8oxoG, **Figure 3A**) on the progression of human pol δ holoenzymes. 8oxoG is one of the most abundant DNA lesions generated by exposure of genomic DNA to reactive oxygen species (ROS) (33). The P/T DNA substrate (Bio-Cy5-P/T-8oxoG, **Figure S1**) contains an 8oxoG 12 nt downstream of the P/T junction (at the 12^th^ dNTP incorporation step, *i*_12_). As observed in **Figure 3B** for an 8oxoG DNA lesion, synthesis of the full-length (62-mer) primer extension product is clearly observed, indicating that human pol δ supports stable incorporation of dNTPs opposite and beyond an 8oxoG (i.e., lesion bypass) during an initial encounter. The observed *P*_i_ values up to, but not including, the 12^th^ dNTP incorporation step (*i*_12_) are identical for the native and 8oxoG P/T DNA substrates (**Figure 3C**). Thus, a downstream 8oxoG DNA does not affect the progression of pol δ holoenzymes towards the lesion. In other words, a downstream 8oxoG lesion does not cause progressing pol δ holoenzymes to prematurely dissociate before the lesion is encountered. Upon encountering an 8oxoG lesion, only 36.4 ± 1.48 % of the progressing pol δ holoenzymes bypass the lesion prior to dissociation (**Table S1**, Probability of bypass x 100%), which is significantly less than that observed for bypass of native G (97.1 ± 0.0606%) in the same sequence context. The reduced efficiency (37.5 ± 1.52 %) of 8oxoG bypass (**Figure 3D**) is primarily due to reduced extension efficiency (43.7 ± 1.45 %) following moderately efficient insertion opposite 8oxoG (85.7 ± 0.646 %). Immediately following lesion bypass of 8oxoG (*i*_12_ and *i*_13_), the observed *P*_i_ values (from *i*_14_ to *i*_33_) are restored to those observed for the native G template (**Figure 3B**). Thus, after bypass of an 8oxoG, the offending lesion does not affect the progression of pol δ holoenzymes that continue downstream. In other words, an 8oxoG lesion that has been bypassed does not cause pol δ holoenzymes that continue downstream to prematurely dissociate downstream before the end of the template is reached. Altogether, this indicates that an 8oxoG lesion only promotes dissociation of pol δ during lesion bypass (i.e., insertion and extension).

**Figure 3.**
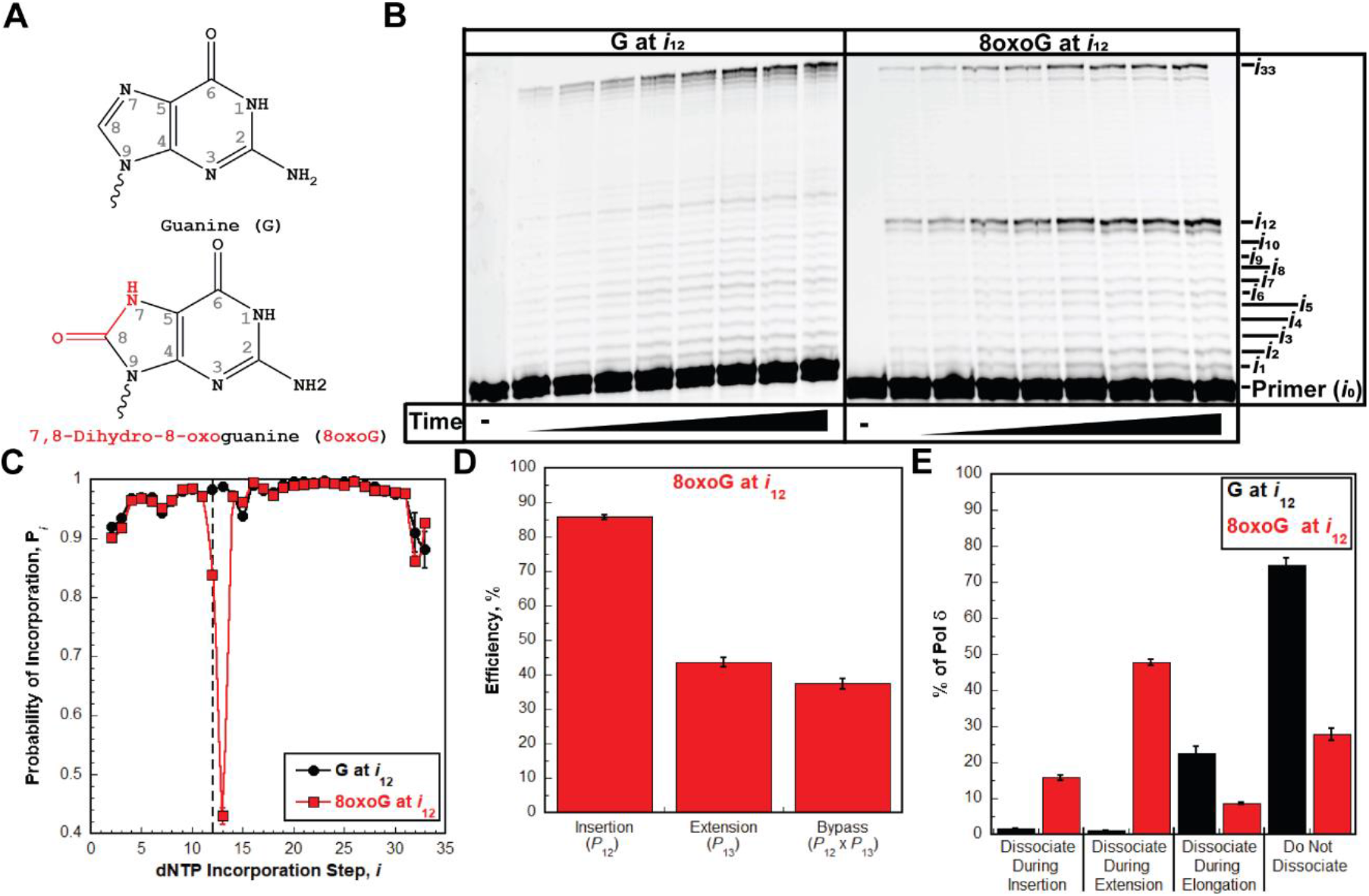
Pol δ holoenzymes encountering an 8oxoG lesion downstream of a P/T junction. The progression of human pol δ holoenzymes was monitored on a P/T DNA substrate (Bio-Cy5-P/T-8oxoG, **Figure S1**) that contains an 8oxoG 12 nt downstream of the P/T junction (at the 12^th^ dNTP incorporation step, *i*_12_). (**A**) Structure of 8oxoG. 8oxoG (*Bottom*) is generated from G (*Top*) through the introduction of an oxo group on the carbon at position 8 (8) and the addition of a hydrogen to the nitrogen at position 7 (7). These modifications are highlighted in red on the structure of 8oxoG. (**B**) 16% denaturing sequencing gel of the primer extension products. The incorporation step (*i*) for certain primer extension products (*i*_*1*_ to *i*_10_, *i*_12_, and *i*_33_) is indicated on the far right. Shown on the left and the right are representative gels of primer extension by pol δ holoenzymes on the native Bio-Cy5-P/T (“G at *i*_12_”) and the Bio-Cy5-P/T-8oxoG (“8oxoG at *i*_12_”) DNA substrates, respectively. (**C**) Processivity of pol δ holoenzymes. *P*_i_ values observed for the native Bio-Cy5-P/T (“G at *i*_12_”) and the Bio-Cy5-P/T-8oxoG (“8oxoG at *i*_12_”) DNA substrates are shown in black and red, respectively, and plotted as a function of the dNTP incorporation step, *i*. Each data point represents the average ± S.E.M. of 3 independent experiments. Error bars are present for all data points but may be smaller than the data point. Data is fit to an interpolation only for observation. Dashed line indicates dNTP incorporation step for insertion (*i*_12_). (**D**) Efficiency of replicating 8oxoG. The efficiencies for insertion, extension, and bypass (insertion and extension) are calculated as described in *Experimental Procedures* and plotted as percentages. The *P*_i_ value(s) from which each efficiency is derived from is indicated below the respective efficiency. Each column represents the average ± S.E.M. of 3 independent experiments. Values for each parameter are also reported in **Table S1**. (**B**). Dissociation of pol δ holoenzymes after encountering an 8oxoG lesion at *i*_12_. The distribution of pol δ dissociation events observed for the native Bio-Cy5-P/T (“G at *i*_12_”) and the Bio-Cy5-P/T-8oxoG (“8oxoG at *i*_12_”) DNA substrates are indicated in black and red, respectively. Error bars reflect the standard error of the mean of 3 independent experiments.

Next, we further assessed the effects of an 8oxoG on the progression of pol δ holoenzymes that encounter the lesion. To do so, we calculated and directly compared the distributions of pol δ dissociation events that occur after an 8oxoG or a native G is encountered at *i*_12_ (**Figure 3E)**. For pol δ holoenzymes that encounter a native G at *i*_12_ (“G at *i*_12_ in **Figure 3E**), the vast majority of the associated pol δ (74.7 ± 2.04 %) does not dissociate at all before reaching the end of the template (*i*_32_ and *i*_33_), as expected. Of the dissociation events that do occur, nearly all are observed during elongation. For pol δ holoenzymes that encounter an 8oxoG at *i*_12_ (“8oxoG at *i*_12_ in **Figure 3E**), only 15.8 ± 0.634 % of pol δ dissociates during insertion, indicating that nearly all (84.2 ± 0.634%) 8oxoG lesions encountered by progressing pol δ holoenzymes are replicated by pol δ. Dissociation of pol δ is most prevalent during extension (47.8 ± 0.848%) but a significant portion of pol δ (27.8 ± 1.62 %) does not dissociate at all before reaching the end of the template.

The intrinsic 3′→5′ exonuclease (i.e., proofreading) activity of human pol δ may affect 8oxoG bypass. To examine this possibility, we repeated the assays and analyses described above with exonuclease-deficient human pol δ. Under the conditions of the assay, where only initial binding encounters of pol δ are monitored, any observed proofreading occurs *intrinsically* (as opposed to *extrinsically*) because a given dNTP incorporation and the subsequent proofreading of that dNTP incorporation are not separated by a dissociation event. If lesion bypass is restricted by the proofreading activity of pol δ, then disabling this activity will increase efficiency of lesion bypass by promoting insertion, extension, or both activities, resulting in an increase in the percentage of pol δ that does not dissociate and a shift of the observed dissociation events towards elongation (34-36). Conversely, if lesion bypass is promoted by the proofreading activity of pol δ, then disabling this activity will decrease the efficiency of lesion bypass by prohibiting insertion, extension, or both activities, resulting in a decrease in the percentage of pol δ that does not dissociate and a shift in the observed dissociation events towards insertion and/or extension. As observed in **Figure 4A** and **Table S1**, disabling the proofreading activity of human pol δ marginally decreases the bypass efficiency (by 8.91 ± 1.84 %) by slightly decreasing the insertion efficiency (by 9.88 ± 0.906 %) and the extension efficiency (by 6.01 ± 2.07 %). This results in a decrease in the percentage of pol δ that does not dissociate and a shift in dissociation events to insertion (**Figure 4B**). Altogether, the studies described above suggest that human pol δ holoenzymes are remarkably efficient at replicating 8oxoG in a lagging strand template (insertion efficiency = 85.7 ± 0.646 %) and that 8oxoG only promotes dissociation of pol δ during lesion bypass; 8oxoG does not affect the progression of pol δ holoenzymes towards the lesion (before lesion bypass) or 2 nt beyond the lesion (after lesion bypass). Furthermore, the 3′→5′ exonuclease activity of human pol δ marginally promotes 8oxoG bypass by proofreading insertion opposite the lesion and potentially extension beyond the lesion. Under the conditions of the assay, it cannot be discerned whether the contribution of proofreading to extension is due to proofreading insertion of an incorrect dNTP opposite 8oxoG (i.e., mismatch) to promote extension or to proofreading extension to stabilize dNTP incorporation 1 nt downstream of the lesion. Next, we repeated these assays to analyze the effects of another prominent oxidative DNA lesion, 5,6-dihydroxy-5,6-dihydrothymine, i.e. thymine glycol (Tg, **Figure 5A**), on the progression of human pol δ holoenzymes (37). Thymine is the most oxidized DNA nucleobase and thymine glycol is the most common oxidation product of thymine (37).

**Figure 4.**
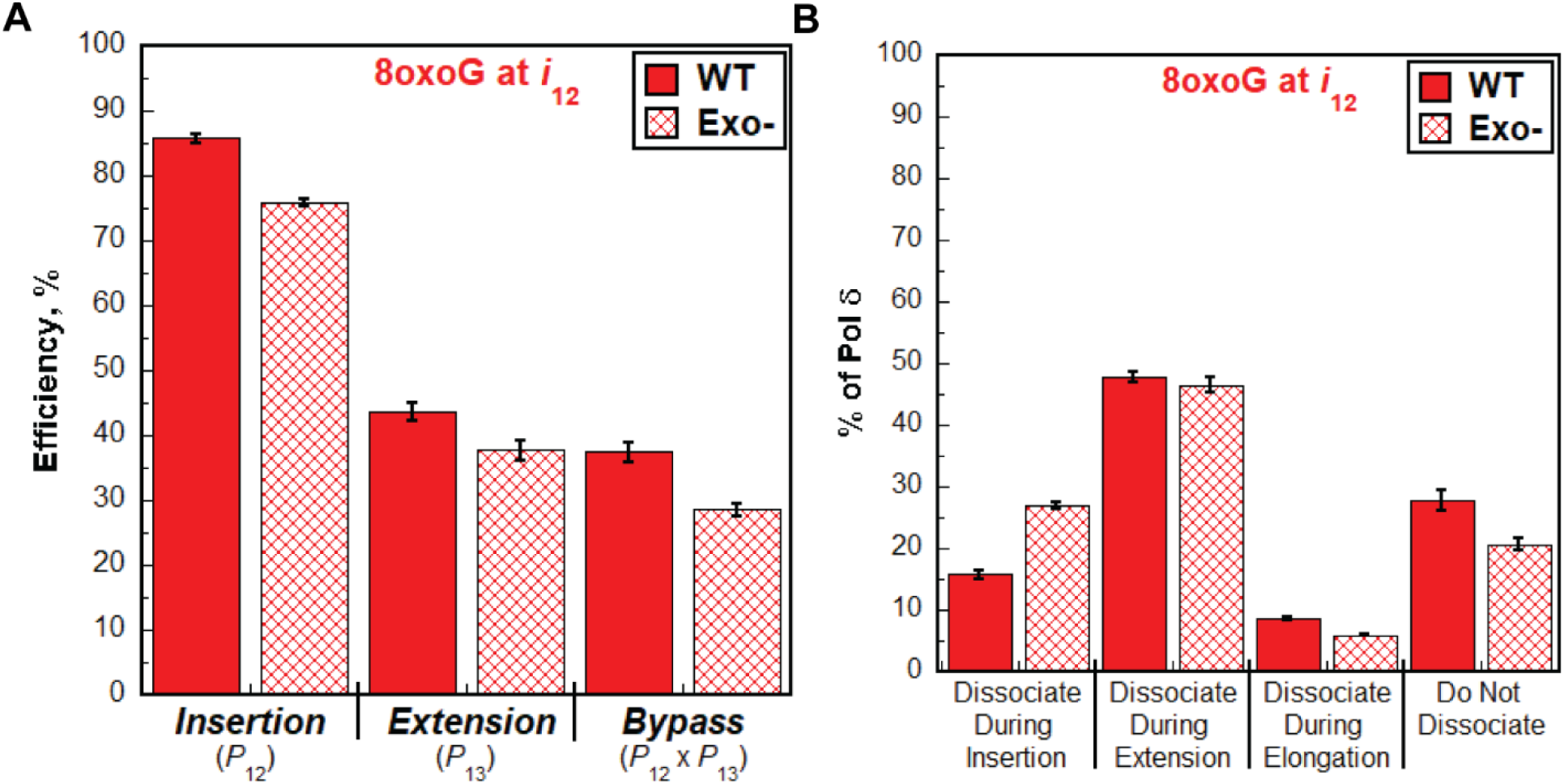
Effect of proofreading on bypass of 8oxoG by pol δ holoenzymes. (**A**) Efficiency of replicating 8oxoG. The efficiencies for dNTP incorporation opposite 8oxoG (i.e., insertion), 1 nt downstream of 8oxoG (i.e., extension), and bypass for wild-type (WT) and exonuclease-deficient (Exo-) pol δ holoenzymes are plotted as percentages. The *P*_i_ value(s) from which each efficiency is derived from is indicated below the respective efficiency. Each column represents the average ± S.E.M. of 3 independent experiments. Values for each parameter are also reported in **Table S1**. (**B**) Dissociation of pol δ holoenzymes after encountering an 8oxoG lesion at *i*_12_. The distribution of dissociation events observed for the Bio-Cy5-P/T-8oxoG DNA substrate with wild-type (WT) and exonuclease-deficient (Exo-) pol δ holoenzymes are plotted. Error bars reflect the standard error of the mean of 3 independent experiments.

**Figure 5.**
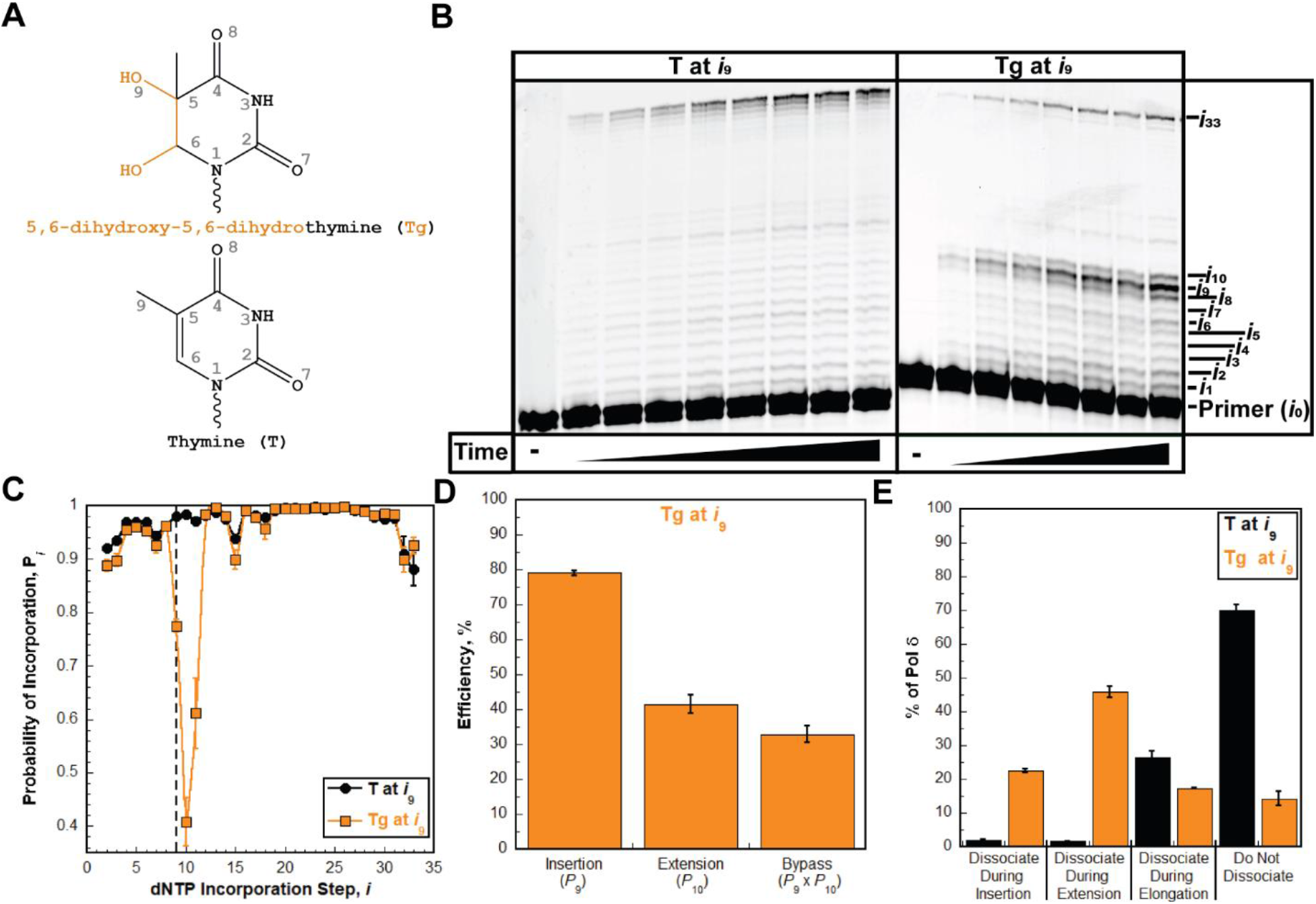
Pol δ holoenzymes encountering a Tg lesion downstream of a P/T junction. The progression of human pol δ holoenzymes was monitored on a P/T DNA substrate (Bio-Cy5-P/T-Tg, **Figure S1**) that contains a Tg 9 nt downstream of the P/T junction (at the 9^th^ dNTP incorporation step, *i*_9_). (**A**) Structure of Tg. Tg (*Bottom*) is generated from T (*Top*) through the addition of hydroxyl groups on the carbons at position 5 (5) and position 6 (6) of the ring. This results in a loss of aromaticity and conversion from planar to nonplanar. These modifications are highlighted in orange on the structure of Tg. (**B**) 16% denaturing sequencing gel of the primer extension products. The incorporation step (*i*) for certain primer extension products (*i*_*1*_ to *i*_10_ and *i*_33_) is indicated on the far right. Shown on the left and the right are representative gels of primer extension by pol δ holoenzymes on the native Bio-Cy5-P/T (“T at *i*_9_”) and the Bio-Cy5-P/T-Tg (“Tg at *i*_9_”) DNA substrates, respectively. (**C**) Processivity of pol δ holoenzymes. *P*_i_ values observed for the native Bio-Cy5-P/T (“T at *i*_9_”) and the Bio-Cy5-P/T-Tg (“Tg at *i*_9_”) DNA substrates are shown in black and orange, respectively, and plotted as a function of the dNTP incorporation step, *i*. Each data point represents the average ± S.E.M. of 3 independent experiments. Error bars are present for all data points but may be smaller than the data point. Data is fit to an interpolation only for observation. Dashed line indicates dNTP incorporation step for insertion (*i*_9_). (**D**) Efficiency of replicating Tg. The efficiencies for dNTP incorporation opposite Tg (i.e., insertion), 1 nt downstream of Tg (i.e., extension), bypass (insertion and extension), and elongation are calculated as described in *Experimental Procedures* and plotted as percentages. The *P*_i_ value(s) from which each efficiency is derived from is indicated below the respective efficiency. Each column represents the average ± S.E.M. of 3 independent experiments. Values for each parameter are also reported in **Table S2**. (**E**). Dissociation of pol δ holoenzymes after encountering a Tg lesion. The distribution of dissociation events observed for the native Bio-Cy5-P/T (“T at *i*_9_”) and the Bio-Cy5-P/T-Tg (“Tg at *i*_9_”) DNA substrates are indicated by black and orange, respectively. Error bars reflect the standard error of the mean of 3 independent experiments.

The P/T DNA substrate (Bio-Cy5-P/T-Tg, **Figure S1**) contains a Tg 9 nt downstream of the P/T junction (at the 9^th^ dNTP incorporation step, *i*_9_). As observed in **Figure 5B**, synthesis of the full-length (62-mer) primer extension product is clearly observed, indicating that human pol δ supports lesion bypass of a Tg during an initial encounter. The observed *P*_i_ values up to, but not including, the 9^th^ dNTP incorporation step (*i*_9_) are nearly identical for the native and Tg P/T DNA substrates (**Figure 5C**). Thus, a downstream Tg DNA lesion does not significantly affect, if at all, the progression of pol δ holoenzymes towards the lesion. Upon encountering a Tg lesion, only 31.6 ± 2.29 % of replicating pol δ holoenzymes bypass the lesion prior to dissociation, which is significantly less than that observed for bypass of native T (96.3 ± 0.178%) in the same sequence context (**Table 2**). The reduced efficiency (32.9 ± 2.38 %) of Tg bypass (**Figure 5D**) is primarily due to reduced extension efficiency (41.5 ± 2.70 %) following moderately efficient insertion opposite Tg (79.1 ± 0.674 %). Immediately following lesion bypass of Tg, native *P*_*i*_ values are not restored until 3 nt beyond the lesion (at the 12^th^ dNTP incorporation step, **Figure 5C**). Thus, after bypass of a Tg, the offending lesion promotes dissociation of pol δ holoenzymes that continue downstream. Altogether, this indicates that a Tg lesion promotes dissociation of pol δ during insertion, extension, and the 1^st^ dNTP incorporation step of elongation (i.e., *i*_11_). However, only 22.5 ± 0.656 % of pol δ dissociates during insertion (**Figure 5E**), indicating that nearly all (77.5 ± 0.656 %) Tg lesions encountered by progressing pol δ holoenzymes are replicated by this DNA polymerase. Dissociation of pol δ is most prevalent during extension (45.9 ± 1.68 %) but a significant portion (17.4 ± 0.267 %) dissociates during elongation, primarily at *i*_11_. Furthermore, 14.3 ± 2.12 % of pol δ does not dissociate at all before reaching the end of the template.

As observed in **Figure 6A**, disabling the 3′→5′ exonuclease activity of human pol δ significantly decreases the bypass efficiency (by 22.9 ± 2.49 %) primarily by decreasing the extension efficiency (by 28.2 ± 2.93 %); the insertion efficiency is only reduced by 3.73 ± 1.32 %. This results in a significant decrease in the percentage of pol δ that does not dissociate and a shift in dissociation events primarily to extension (**Figure 6B**). Altogether, the studies described above suggest that human pol δ holoenzymes are remarkably efficient at replicating Tg in a lagging strand template (insertion efficiency = 79.1 ± 0.674 %) and that Tg does not affect the progression of pol δ holoenzymes towards the lesion (before lesion bypass) but promotes dissociation of pol δ during and after lesion bypass. Furthermore, the 3′→5′ exonuclease activity of human pol δ significantly promotes Tg bypass primarily by promoting extension. Again, under the conditions of the assay, it cannot be discerned whether the significant contribution of proofreading to extension is due to proofreading insertion of an incorrect dNTP opposite a Tg (i.e., mismatch) to promote extension or proofreading extension to stabilize dNTP incorporation 1 nt downstream of the lesion. Next, we examined the effects of prominent alkylative DNA lesions on the progression of human pol δ holoenzymes.

**Figure 6.**
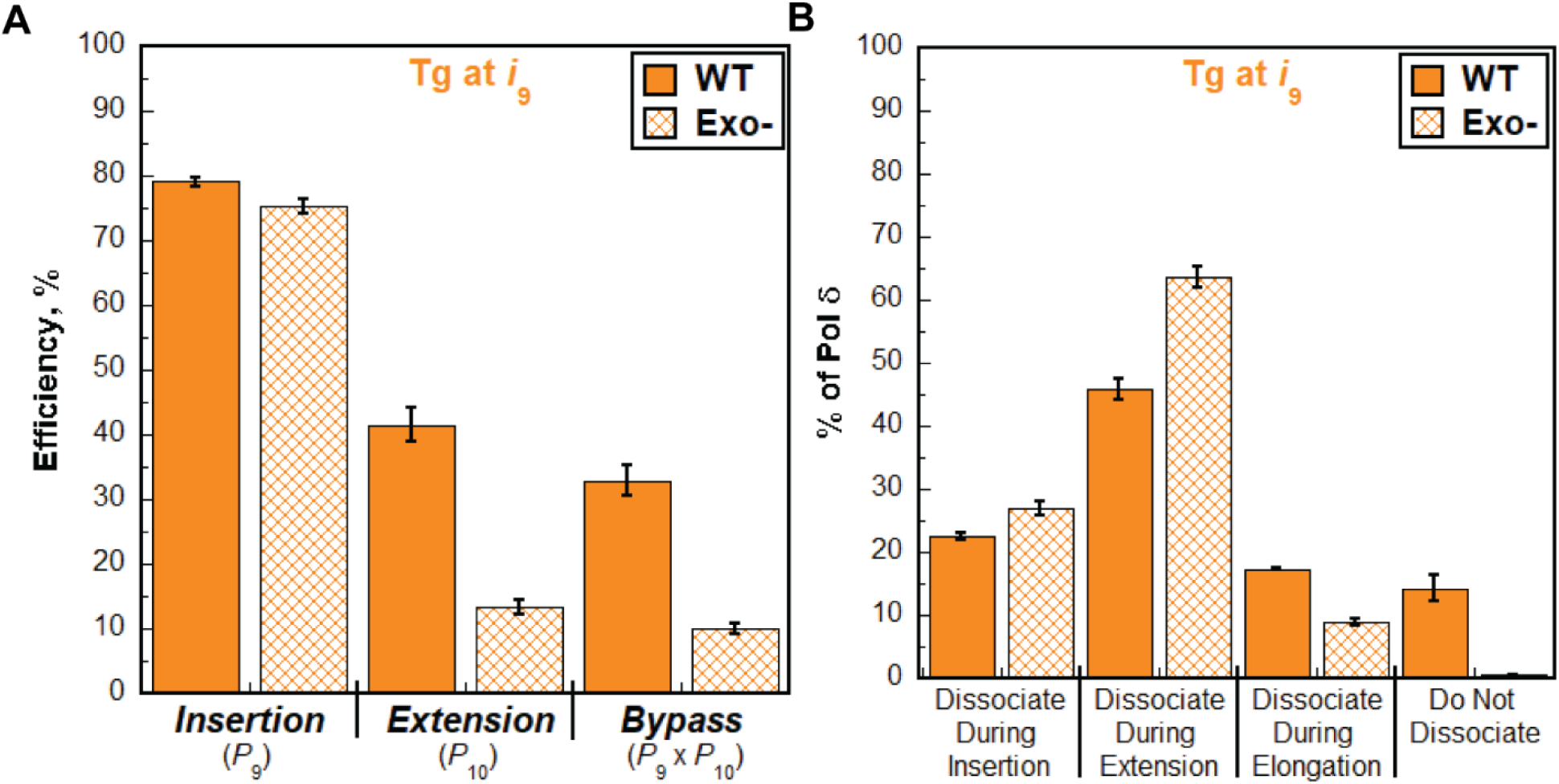
Effect of proofreading on bypass of Tg by pol δ holoenzymes. (**A**) Efficiency of replicating Tg. The efficiencies for dNTP incorporation opposite Tg (i.e., insertion), 1 nt downstream of Tg (i.e., extension), and bypass for wild-type (WT) and exonuclease-deficient (Exo-) pol δ holoenzymes are plotted as percentages. The *P*_i_ value(s) from which each efficiency is derived from is indicated below the respective efficiency. Each column represents the average ± S.E.M. of 3 independent experiments. Values for each parameter are also reported in **Table S2. (E)** Dissociation of pol δ holoenzymes after encountering an Tg lesion. The distribution of dissociation events observed for the Bio-Cy5-P/T-Tg DNA substrate with wild-type (WT) and exonuclease-deficient (Exo-) pol δ holoenzymes are plotted. Error bars reflect the standard error of the mean of 3 independent experiments.

### The effect of alkylative DNA lesions on the progression of human pol δ holoenzymes

First, we examined the effect of *O*^*6*^-Methylguanine (O6MeG, **Figure 7A**) on the progression of human pol δ holoenzymes. O6MeG is a prominent DNA lesion generated by exposure of genomic DNA to methylating agents, such as the antitumor agents dacarbazine, streptozotocin, procarbazine and temozolomide (38). The P/T DNA substrate (Bio-Cy5-P/T-O6MeG, **Figure S1**) contains an O6MeG 12 nt downstream of the P/T junction (at the 12^th^ dNTP incorporation step, *i*_12_). As observed in **Figure 7B**, synthesis of the full-length (62-mer) primer extension product is clearly observed, indicating that human pol δ supports stable incorporation of dNTPs opposite and beyond an O6MeG (i.e., lesion bypass) during an initial encounter. The observed *P*_i_ values up to, but not including, the 12^th^ dNTP incorporation step (*i*_12_) are essentially identical for the native and O6MeG P/T DNA substrates (**Figure 7C**). Thus, a downstream O6MeG DNA lesion does not affect the progression of pol δ holoenzymes towards the lesion. Upon encountering an O6MeG lesion, 66.8 ± 0.60 % of replicating pol δ holoenzymes bypass the lesion prior to dissociation, which is reduced compared to that observed for bypass of native G (97.1 ± 0.0606%) in the same sequence context (**Table S3**). The marginally reduced efficiency of O6MeG bypass (68.8 ± 0.62 %, **Figure 7D, Table S3**) is due to moderate reductions in both the insertion (79.6 ± 0.617 %) and extension efficiencies (86.5 ± 0.25 %). Immediately following O6MeG bypass, native *P*_i_ values are restored (at the 14^th^ dNTP incorporation step, *i*_14_, **Figure 7C**), indicating that, after bypass of an O6MeG, the offending lesion does not affect the progression of pol δ holoenzymes that continue downstream. Altogether, this indicates that an O6MeG lesion only promotes dissociation of pol δ during lesion bypass (i.e., insertion and extension). However, only 21.8 ± 0.605 % of pol δ dissociates during insertion (**Figure 7E**), indicating that nearly all (78.2 ± 0.605 %) O6MeG lesions encountered by progressing pol δ holoenzymes are replicated by pol δ. Dissociation of pol δ is less prevalent during extension (11.4 ± 0.188 %) and elongation (17.5 ± 0.414 %) and, surprisingly, half of pol δ (49.3 ± 0.87 %) does not dissociate at all before reaching the end of the template.

**Figure 7.**
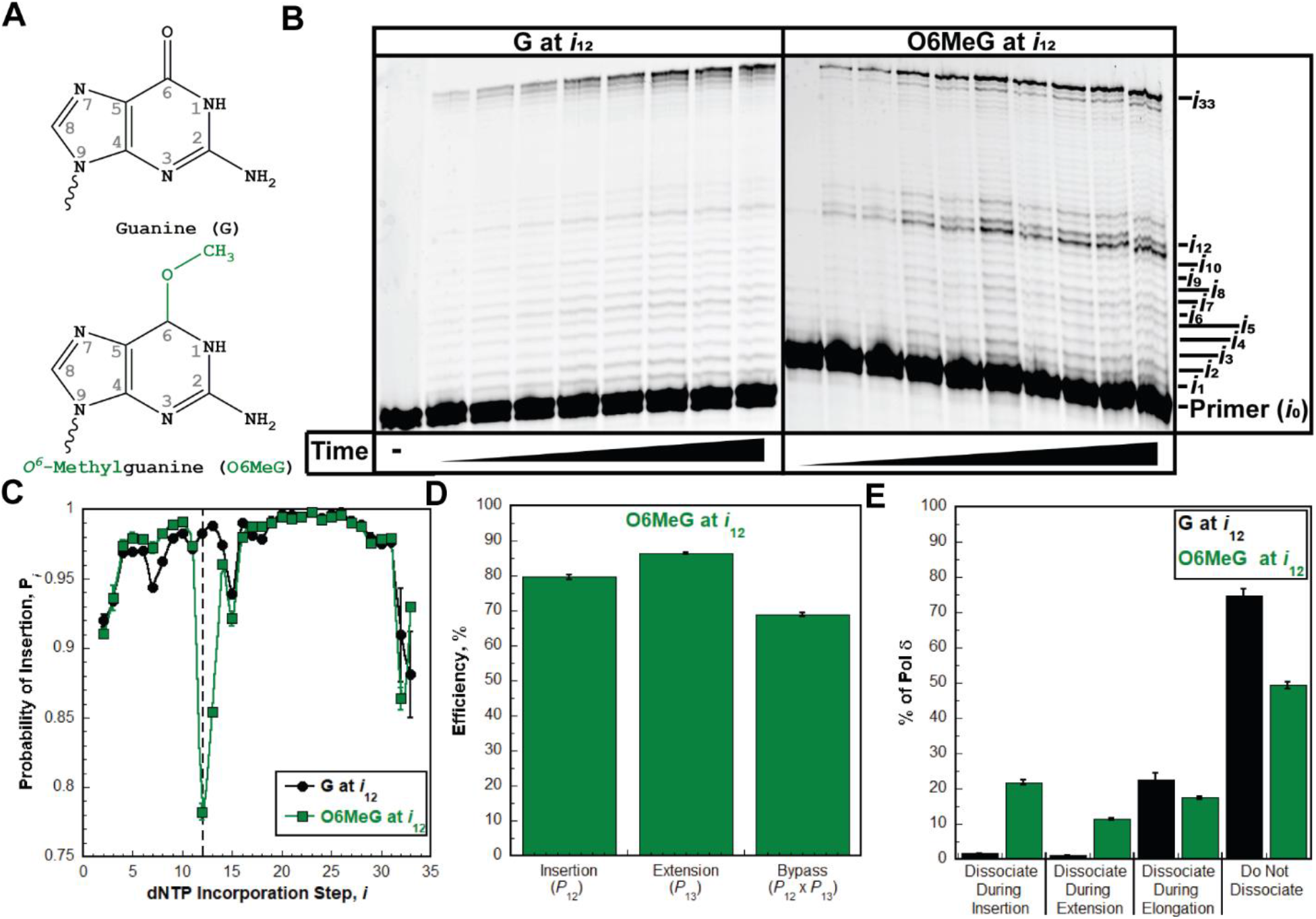
Pol δ holoenzymes encountering an O6MeG lesion downstream of a P/T junction. The progression of human pol δ holoenzymes was monitored on a P/T DNA substrate (Bio-Cy5-P/T-O6MeG, **Figure S1**) that contains an O6MeG 12 nt downstream of the P/T junction (at the 12^th^ dNTP incorporation step, *i*_12_). (**A**) Structure of O6MeG. O6MeG (*Bottom*) is generated from G (*Top*) through the addition of a methyl group on the oxygen of the carbonyl group at position 6 (6) of the ring. These modifications are highlighted in green on the structure of O6MeG. (**B**) 16% denaturing sequencing gel of the primer extension products. The incorporation step (*i*) for certain primer extension products (*i*_*1*_ to *i*_12_ and *i*_33_) is indicated on the far right. Shown on the left and the right are representative gels of primer extension by pol δ holoenzymes on the native Bio-Cy5-P/T (“G at *i*_12_”) and the Bio-Cy5-P/T-O6MeG (“O6MeG at *i*_12_”) DNA substrates, respectively. (**C**) Processivity of pol δ holoenzymes. *P*_i_ values observed for the native Bio-Cy5-P/T (“G at *i*_12_”) and the Bio-Cy5-P/T-O6MeG (“O6MeG at *i*_12_”) DNA substrates are shown in black and green, respectively, and plotted as a function of the dNTP incorporation step, *i*. Each data point represents the average ± S.E.M. of 3 independent experiments. Error bars are present for all data points but may be smaller than the data point. Data is fit to an interpolation only for observation. Dashed line indicates dNTP incorporation step for insertion (*i*_12_). (**D**) Efficiency of replicating O6MeG. The efficiencies for dNTP incorporation opposite O6MeG (i.e., insertion), 1 nt downstream of O6MeG (i.e., extension), and bypass (insertion and extension) are calculated as described in *Experimental Procedures* and plotted as percentages. The *P*_i_ value(s) from which each efficiency is derived from is indicated below the respective efficiency. Each column represents the average ± S.E.M. of 3 independent experiments. Values for each parameter are also reported in **Table S3**. (**E**). Dissociation of pol δ holoenzymes after encountering an O6MeG lesion. The distribution of dissociation events observed for the native Bio-Cy5-P/T (“G at *i*_12_”) and the Bio-Cy5-P/T-O6MeG (“O6MeG at *i*_12_”) DNA substrates are indicated by black and green, respectively. Error bars reflect the standard error of the mean of 3 independent experiments.

As observed in **Figure 8A**, disabling the 3′→5′ exonuclease activity of human pol δ significantly decreases the bypass efficiency for O6MeG (by 33.0 ± 1.27 %) by slightly decreasing the insertion efficiency (by 12.6 ± 1.04 %) and significantly decreasing the extension efficiency (by 33.1 ± 1.02 %). This results in a drastic decrease in the percentage of pol δ that does not dissociate (**Figure 8B**) and a significant increase in the percentage of pol δ that dissociates during insertion (21.8 ± 0.605% to 35.4 ± 0.796%), extension (11.4 ± 0.188% to 31.3 ± 0.226 %), as well as elongation (17.5 ± 0.414% to 29.2 ± 0.803%). Altogether, these studies suggest that human pol δ holoenzymes are remarkably efficient at replicating O6MeG in a lagging strand template (insertion efficiency = 79.6 ± 0.617 %) and that O6MeG only promotes dissociation of pol δ during lesion bypass; O6MeG does not affect the progression of pol δ holoenzymes towards the lesion (before lesion bypass) or 2 nt beyond the lesion (after lesion bypass). Furthermore, the 3′→5′ exonuclease activity of human pol δ significantly promotes O6MeG bypass by proofreading insertion opposite the lesion and potentially extension beyond the lesion. Again, under the conditions of the assay, it cannot be discerned whether the significant contribution of proofreading to extension is due to proofreading insertion of an incorrect dNTP opposite O6MeG (i.e., mismatch) to promote extension or proofreading extension to stabilize dNTP incorporation 1 nt downstream of the lesion. Finally, we repeated these assays to analyze the effects of another prominent alkylative DNA lesion, *1,N*^*6*^-ethenoadenine, i.e. ethenoadenine (εA, **Figure 9A**), on the progression of human pol δ holoenzymes. εA is a prominent alkylation product of adenine that is generated by exposure of genomic DNA to vinyl chloride, an industrial pollutant, or lipid peroxidation byproducts associated with inflammation and metabolism (39).

**Figure 8.**
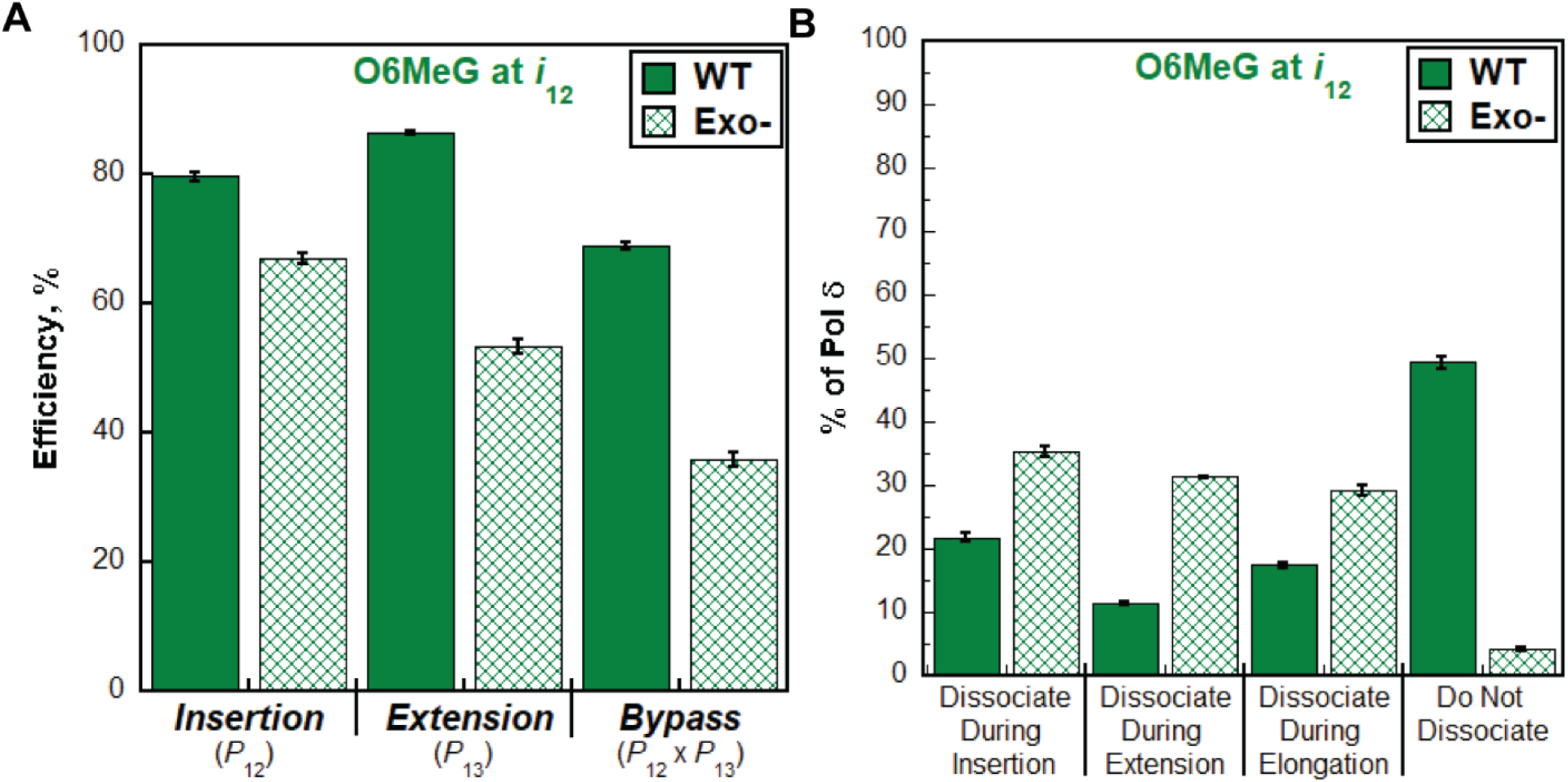
Effect of proofreading on bypass of O6MeG by pol δ holoenzymes. (**A**) Efficiency of replicating O6MeG. The efficiencies for dNTP incorporation opposite O6MeG (i.e., insertion), 1 nt downstream of O6MeG (i.e., extension) and bypass for wild-type (WT) and exonuclease-deficient (Exo-) pol δ holoenzymes are plotted as percentages. The *P*_i_ value(s) from which each efficiency is derived from is indicated below the respective efficiency. Each column represents the average ± S.E.M. of 3 independent experiments. Values for each parameter are also reported in **Table S3**. (**B**) Dissociation of pol δ holoenzymes after encountering an O6MeG lesion. The distribution of dissociation events observed for the Bio-Cy5-P/T-O6MeG DNA substrate with wild-type (WT) and exonuclease-deficient (Exo-) pol δ holoenzymes are plotted. Error bars reflect the standard error of the mean of 3 independent experiments.

**Figure 9.**
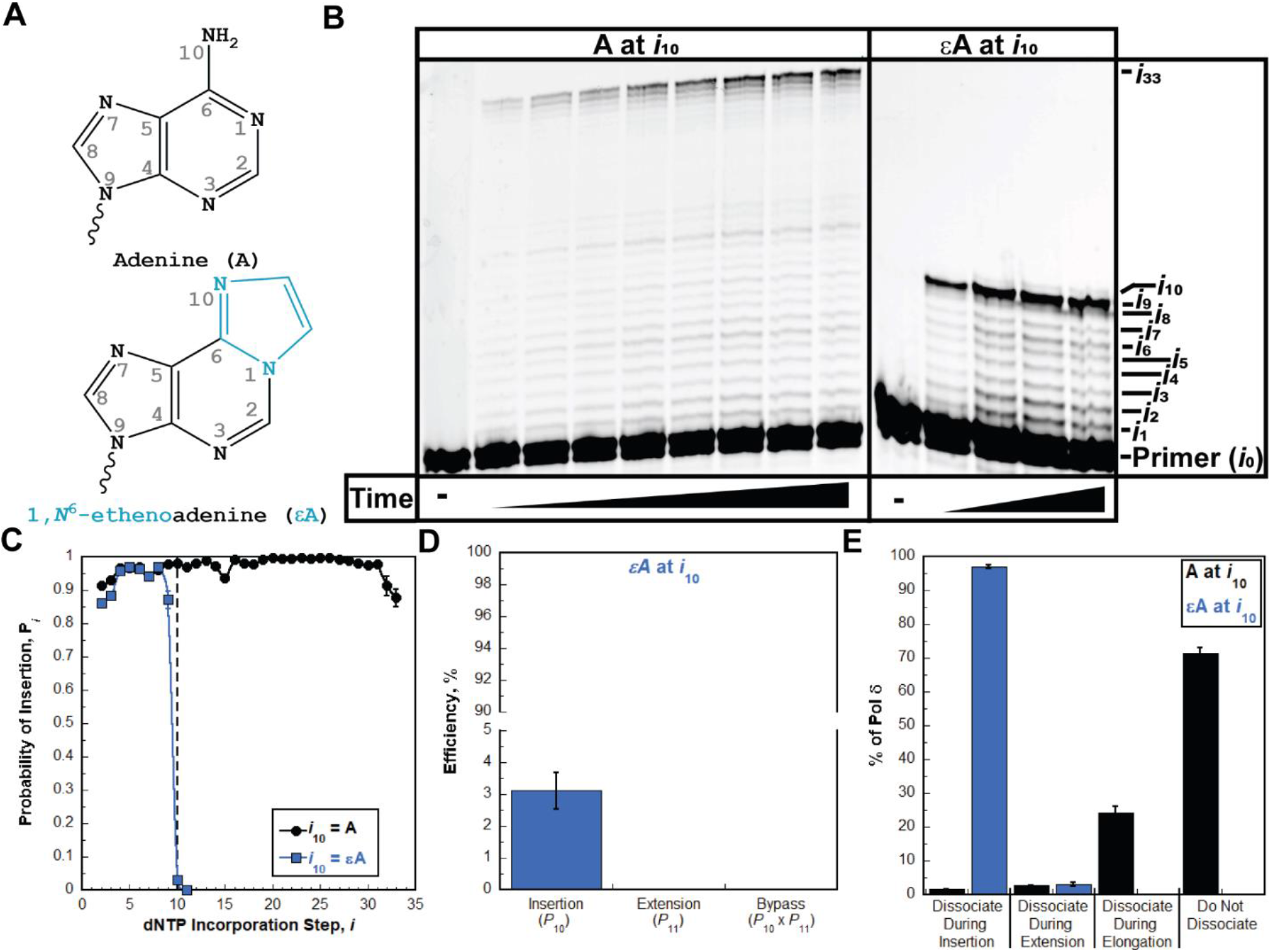
Pol δ holoenzymes encountering an εA lesion downstream of a P/T junction. The progression of human pol δ holoenzymes was monitored on a P/T DNA substrate (Bio-Cy5-P/T-εA, **Figure S1**) that contains an εA 10 nt downstream of the P/T junction (at the 10^th^ dNTP incorporation step, *i*_10_). (**A**) Structure of εA. εA (*Bottom*) is generated from A (*Top*) through the attachment of 2 extra carbons in an exocyclic arrangement; 1 carbon is attached to the nitrogen at position 1 (1) and the other is attached to the nitrogen in the amine at position 6 (6) of the ring. These modifications are highlighted in blue on the structure of εA. (**B**) 16% denaturing sequencing gel of the primer extension products. The dNTP incorporation step (*i*) for certain primer extension products (*i*_*1*_ to *i*_10_ and *i*_33_) is indicated on the far right. Shown on the left and the right are representative gels of primer extension by pol δ holoenzymes on the native Bio-Cy5-P/T (“A at *i*_10_”) and the Bio-Cy5-P/T-εA (“εA at *i*_10_”) DNA substrates, respectively. (**C**) Processivity of pol δ holoenzymes. *P*_i_ values observed for the native Bio-Cy5-P/T (“A at *i*_10_”) and the Bio-Cy5-P/T-εA (“εA at *i*_10_”) DNA substrates are shown in black and blue, respectively, and plotted as a function of the dNTP incorporation step, *i*. Each data point represents the average ± S.E.M. of 3 independent experiments. Error bars are present for all data points but may be smaller than the data point. Data is fit to an interpolation only for observation. (**D**) Efficiency of replicating εA. The efficiencies for dNTP incorporation opposite εA (i.e., insertion), 1 nt downstream of εA (i.e., extension) and bypass (insertion and extension) are calculated as described in *Experimental Procedures* and plotted as percentages. The *P*_i_ value(s) from which each efficiency is derived from is indicated below the respective efficiency. Each column represents the average ± S.E.M. of 3 independent experiments. Values for each parameter are also reported in **Table S4**. (**E**). Dissociation of pol δ holoenzymes after encountering an εA lesion. The distribution of dissociation events observed for the native Bio-Cy5-P/T (“X = A”) and the Bio-Cy5-P/T-εA (“X = εA”) DNA substrates are indicated by black and blue, respectively. Error bars reflect the standard error of the mean of 3 independent experiments.

The P/T DNA substrate (Bio-Cy5-P/T-εA, **Figure S1**) contains an εA 10 nt downstream of the P/T junction (at the 10^th^ dNTP incorporation step, *i*_10_). As observed in **Figure 9B**, synthesis past the εA DNA lesion does not occur, indicating that human pol δ does not support lesion bypass of an εA during an initial encounter. Interestingly, the observed *P*_i_ values for the native and εA P/T DNA substrates are nearly identical up to only the 8^th^ dNTP incorporation step (*i*_8_) and then significantly diverge at the 9^th^ dNTP incorporation step (*i* _9_) and beyond (**Figure 9C**). This suggests that a downstream εA DNA lesion may cause some progressing pol δ holoenzymes to prematurely dissociate before the lesion is encountered. Upon encountering an εA lesion, only 3.06 ± 0.558 % of progressing pol δ holoenzymes incorporate a dNTP opposite the lesion prior to dissociation, which is drastically less than that observed for bypass of native A (98.3 ± 0.0848%) in the same sequence context (**Table S4**). Hence, the efficiency for dNTP incorporation opposite an εA lesion is extremely low (3.11 ± 0.568 %, **Figure 9D**) and by far the lowest of all DNA lesions analyzed in the present study. Extension beyond a εA lesion and, hence, bypass and elongation are not observed. Thus, all progressing pol δ holoenzymes that encounter an εA lesion dissociate during insertion or extension (**Figure 9E**). The former accounts for 96.9 ± 0.558 % of all dissociation events. As observed in **Figure 10A**, disabling the 3′→5′ exonuclease activity of human pol δ slightly increased the insertion efficiency, if at all, and did not yield extension. Furthermore, the observed distribution of dissociation events is not visibly altered by disabling the 3′→5′ exonuclease activity of human pol δ (**Figure 10B**). This suggests that the 3′→5′ exonuclease activity of human pol δ does not contribute to dNTP incorporation opposite εA and that insertion may evade intrinsic proofreading by human pol δ. Altogether, these studies suggest that human pol δ holoenzymes are very inefficient at replicating εA in a lagging strand template (insertion efficiency = 3.11 ± 0.568 %%) and that εA promotes dissociation of pol δ during lesion bypass and also as the polymerase approaches the lesion. Furthermore, the 3′→5′ exonuclease activity of human pol δ does not contribute to lesion bypass of εA during initial encounters.

**Figure 10.**
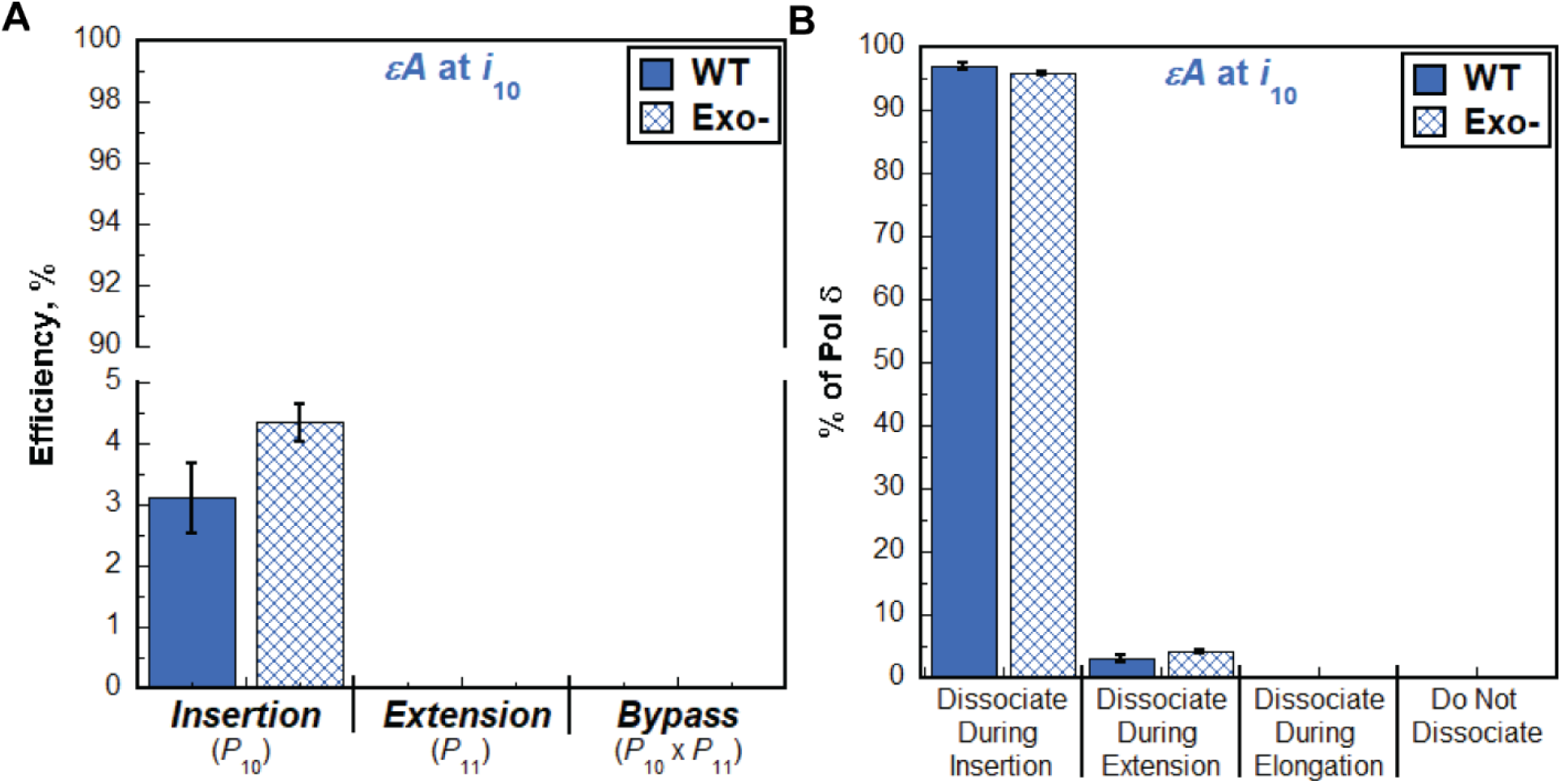
Effect of proofreading on bypass of εA by pol δ holoenzymes. (**A**) Efficiency of replicating εA. The efficiencies for dNTP incorporation opposite εA (i.e., insertion), 1 nt downstream of εA (i.e., extension), and bypass for wild-type (WT) and exonuclease-deficient (Exo-) pol δ holoenzymes are plotted as percentages. The *P*_i_ value(s) from which each efficiency is derived from is indicated below the respective efficiency. Each column represents the average ± S.E.M. of 3 independent experiments. Values for each parameter are also reported in **Table S4**. (**B**) Dissociation of pol δ holoenzymes after encountering an εA lesion. The distribution of dissociation events observed for the Bio-Cy5-P/T-εA DNA substrate with wild-type (WT) and exonuclease-deficient (Exo-) pol δ holoenzymes are plotted. Error bars reflect the standard error of the mean of 3 independent experiments.

## Discussion

In the present study, we re-constituted human lagging strand replication at physiological pH, ionic strength, and dNTP concentrations to quantitatively characterize, at single nucleotide resolution, the initial encounters of pol δ holoenzymes with downstream DNA lesions. In short, a DNA lesion ≥ 9 nt downstream of a P/T junction is encountered only once and only by a progressing pol δ holoenzyme, rather than pol δ alone. To the best of our knowledge, comparable studies on human lagging strand replication have yet to be reported. The results indicate that human pol δ holoenzymes support stable dNTP incorporation opposite and beyond multiple lesions and the extent of these activities depends on the identity of the lesion (**Figure 11A**) and the ability to proofread intrinsically. Surprisingly, the results reveal that human pol δ holoenzymes are remarkably efficient at inserting a dNTP opposite certain DNA lesions, with efficiencies ≥ ∼80 % for 8oxoG, Tg, and O6MeG lesions (**Figure 11A**). Furthermore, the results indicate that after a progressing pol δ holoenzyme encounters a given DNA lesion, subsequent dissociation of pol δ, if it occurs, is not designated to a uninform site relative to the lesion. Rather, pol δ dissociation events are distributed around the lesion. The distributions of pol δ dissociation events are dependent on the identity of the lesion (**Figure 11B**) and the ability to proofread intrinsically. Taken together with previous reports on human pol δ, the results from the present study reveal complexity and heterogeneity in the replication of DNA lesions in lagging strand templates, as discussed in further detail below.

**Figure 11.**
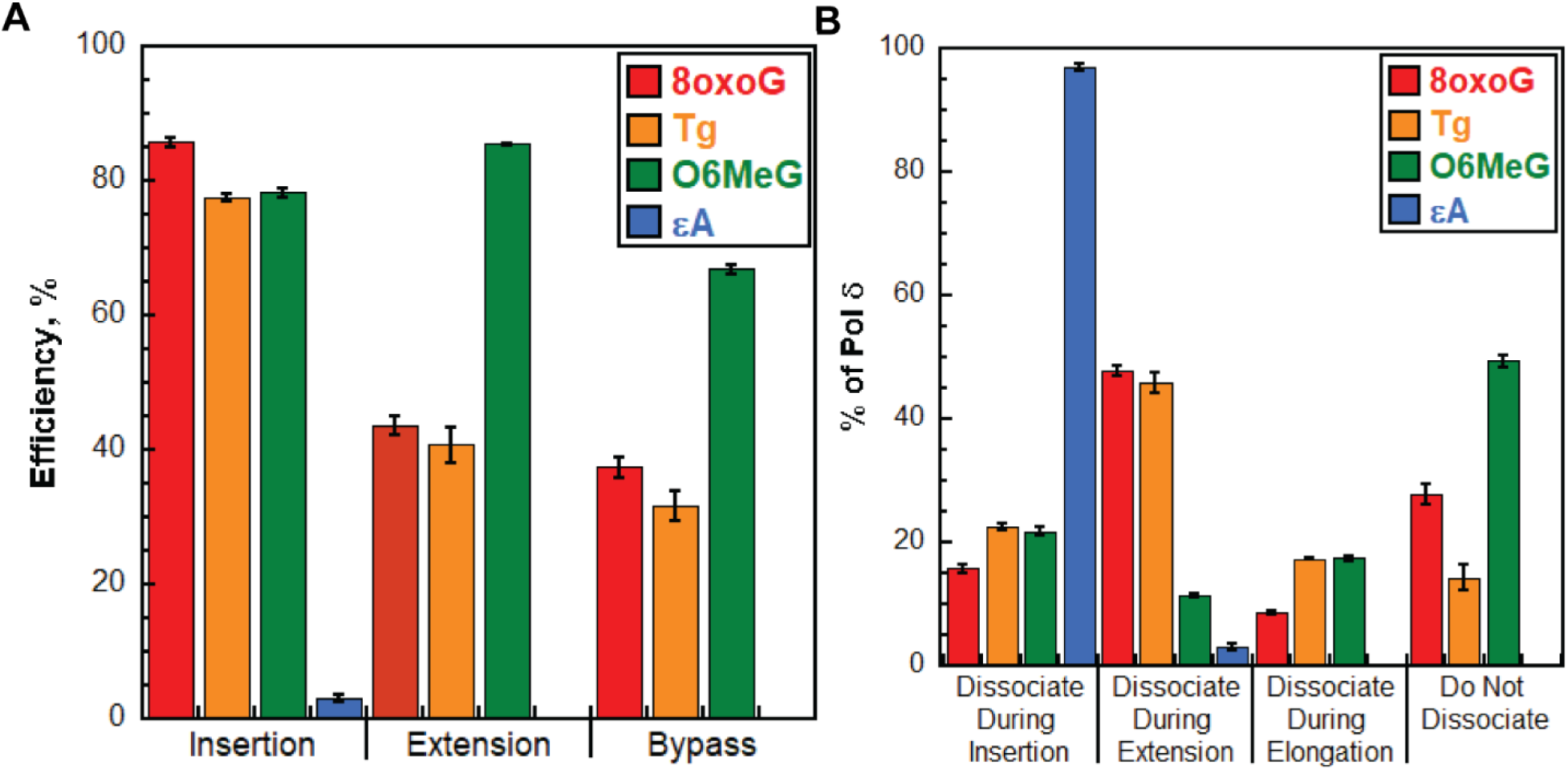
Bypass of DNA lesions by pol δ holoenzymes during initial encounters. (**A**) Efficiencies of replicating DNA lesions. The efficiencies for dNTP incorporation opposite a lesion (i.e., insertion), 1 nt downstream of lesion (i.e., extension) and bypass (insertion and extension) of a lesion for wild type pol δ holoenzymes are plotted. Data is taken from **Figures 3D, 5D, 7D**, and **9D** and is color-coded by DNA lesion. (**B**) Dissociation of pol δ holoenzymes after encountering DNA lesions. The distribution of dissociation events observed for wild type pol δ holoenzymes encountering DNA lesions is plotted. Data is taken from **Figures 3E, 5E, 7E**, and **9E** and is color-coded by DNA lesion.

### 8oxoG lesions in lagging strand templates

Human pol δ holoenzymes are remarkably efficient at inserting a dNTP opposite 8oxoG (insertion efficiency = 85.7 ± 0.646 %, **Figure 11A**) such that 84.2 ± 0.634 % of progressing pol δ holoenzymes that encounter an 8oxoG complete insertion prior to pol δ dissociating (**Figure 11B**). Intrinsic proofreading contributes marginally (9.88 ± 0.906 %) to this unexpectedly high insertion efficiency (**Figure 4A**). Previous studies indicated that, in the presence of PCNA and RPA, human pol δ primarily inserts either the correct dCTP or the incorrect dATP opposite 8oxoG, with the latter accounting for 25 – 40% of all insertion events (8,33,40). In the present study, this equates to ∼21 – 34% of all encounters between progressing pol δ holoenzymes and 8oxoG lesions resulting in 8oxoG:A mismatches and ∼50 – 63% yielding “correct” 8oxoG:C base pairs. Altogether, this suggests that nearly all 8oxoG lesions in lagging strand templates are *initially* replicated by pol δ, rather than a DDT pathway, creating a heterogenous population of nascent DNA that may elicit a variety of downstream responses during DNA replication. For 8oxoG:C base pairs, pol δ faithfully completes insertion and DDT would only be utilized, if at all, to complete extension as dissociation of pol δ is most prevalent at this dNTP incorporation step (**Figure 11B**) and unperturbed progression of pol δ holoenzymes resumes 2 nt downstream of the lesion (**Figure 3C**). For 8oxoG:A mismatches, the mismatched dAMP opposite the 8oxoG lesion must be excised and the 8oxoG accurately “re-replicated” to avoid fixed G:C → T:A transversion mutations. This may occur via multiple pathways that depend on the extent of pol δ holoenzyme progression downstream (3′) of an 8oxoG lesion and/or the activation of DDT pathways. We discuss two possibilities below.

P/T junctions aborted by pol δ at least 6 nt downstream (3′) of a 8oxoG:A mismatch may be recognized and processed by a specialized sub-pathway of base excision repair (BER). Specifically, the human homolog of MutY (MutYH), a monofunctional adenine DNA glycosylase, selectively excises the adenine nucleobase from an 8oxoG:A mismatch, generating an abasic site opposite 8oxoG. Next, apurinic/apyrimidinic endonuclease 1 (APE1) cleaves the phosphodiester bond 5′ to the abasic site, generating a “nick” that permits re-replication of the lesion. Both MutYH and APE1 require ≤ 6 nt of DNA immediately 3′ to their site of action (41-43). In the present study, nearly 40% (38.5 ± 1.75 %) of progressing pol δ holoenzymes that insert a dNTP opposite 8oxoG subsequently extend the nascent DNA at least 6 nt downstream prior to pol δ dissociation and previous studies demonstrated that, in the presence of PCNA and RPA, human pol δ has a slight, albeit undefined, preference for extending 8oxoG:A mismatches compared to 8oxoG:C. Together, this suggests many of the 8oxoG:A mismatches generated by progressing pol δ holoenzymes are subsequently converted by the holoenzymes into substrates for MutYH, and ultimately APE1. The remainder may be extended to sufficient lengths for MutYH and APE1 by pol δ (in subsequent binding encounters) or DDT pathways, if necessary. MutYH protein levels are significantly upregulated in response to oxidative stress (44) and MutYH physically interacts with PCNA and RPA (45), both of which are left behind at P/T junctions aborted by pol δ. Furthermore, APE1 physically interacts with PCNA and MutYH (46). Altogether, this suggests a timely and coordinated pathway for processing of mismatched dAMP opposite 8oxoG during DNA replication. In this scenario, re-replication of 8oxoG occurs within a nick and this may be carried out by a number of DNA polymerases, including pol δ. However, DNA polymerase λ (pol λ) has by far the highest fidelity when replicating 8oxoG, with a mutation frequency less than 0.1 % in the presence of PCNA and RPA (40,47-49). Hence, selection of pol λ would ensure accurate re-replication of 8oxoG. Pol λ levels are significantly upregulated in response to oxidative stress. Furthermore, pol λ physically interacts with MutYH and this interaction is promoted in response to oxidative stress and significantly upregulates chromatin association of pol λ during DNA replication. Finally, pol λ physically interacts with PCNA, which is left behind at P/T junctions aborted by pol δ, and this interaction restricts association of DNA polymerase β (pol β), which replicates 8oxoG with a significantly elevated mutation frequency (25%) compared to pol λ. RPA also contributes to the exclusion of pol β, and potentially other DNA polymerases, by engaging the nick opposite 8oxoG (44,50,51). Altogether, this suggests that pol λ is selected for re-replication of 8oxoG within nicks.

Alternatively, P/T junctions aborted by pol δ shortly downstream (≤ 6 nt) of an 8oxoG:A mismatch may be re-engaged by a factor with 3′ → 5′ exonuclease activity that degrades the nascent DNA to ultimately excise a mismatched dAMP opposite 8oxoG. In the present study, the 3′ → 5′ exonuclease activity of human pol δ significantly promotes bypass of Tg (**Figure 6A, Table 2**) and O6MeG (**Figure 8A, Table 3**) lesions but only had a marginal contribution (< 9%) to the bypass of 8oxoG (**Figure 4A, Table 1**). This reveals that human pol δ holoenzymes are capable of proofreading DNA lesion base pairs but proofreading 8oxoG base pairs is not prevalent. Thus, human pol δ alone is not a strong candidate for excising a mismatched dAMP opposite 8oxoG. An alternative candidate is the Werner syndrome (WRN) protein, which readily excises dNMPs opposite 8oxoG from as far away as 10 nt downstream (3′) of the lesion (52). Previous reports on human DNA replication revealed that WRN physically interacts with PCNA and RPA, both of which are left behind at P/T junctions aborted by pol δ, and, indeed, WRN is recruited to sites of 8oxoG lesions in a DNA replication-dependent manner and suppresses G:C → T:A transversion mutations at 8oxoG lesions *in vivo*. Furthermore, WRN physically interacts with pol δ and the WRN•pol δ complex has enhanced 3′ → 5′ exonuclease activity compared to either component on its own (53-61). Thus, WRN may be recruited to aborted P/T junctions at or downstream (3′) of 8oxoG:A mismatches and directly excise dAMP or facilitate excision by pol δ. In this scenario, re-replication of 8oxoG initiates from a “standing start” on a P/T junction that is encircled by PCNA and directly abuts 8oxoG and a 5′ ssDNA overhang engaged by at least 1 RPA. Nearly all DNA polymerases are tenable to this scenario, pol δ in particular. Hence, in addition to the oxidative stress-induced upregulation of pol λ, complimentary mechanisms may be required to ensure accurate re-replication of 8oxoG lesions in this MutYH-independent scenario. This is currently under investigation.

### Tg lesions in lagging strand templates

Similar to 8oxoG, human pol δ holoenzymes are remarkably efficient at inserting a dNTP opposite Tg (insertion efficiency = 79.1 ± 0.674 %, **Figure 11A**) such that 77.5 ± 0.676 % of progressing pol δ holoenzymes that encounter a Tg complete insertion prior to pol δ dissociating (**Figure 11B**). Furthermore, similar to 8oxoG, intrinsic proofreading slightly contributes (3.73 ± 1.32 %) to this unexpectedly high insertion efficiency (**Figure 6A**). To the best of our knowledge, direct studies on replication of Tg lesions by human pol δ have yet to be reported and, hence, the fidelity of human pol δ in replicating Tg lesions is unknown. However, human DNA polymerase α (pol α), a B-family DNA polymerase like pol δ, exclusively inserts dAMP opposite Tg lesions (62). Furthermore, the DNA polymerase active sites of pol δ’s from *S. cerevisiae* and human are structurally conserved and a recent report demonstrated that *S. cerevisiae* pol δ exclusively inserts dAMP opposite Tg lesions in the presence of PCNA and RPA (63-65). Altogether, this suggests that the overwhelming majority Tg lesions in lagging strand templates are *initially* replicated by pol δ, rather than a DDT pathway, and that replication of Tg lesions by pol δ is error-free. Accordingly, DDT is primarily utilized, if at all, to complete extension and/or the 1^st^ dNTP incorporation of elongation as dissociation of pol δ is most prevalent during these dNTP incorporation steps and unperturbed progression of pol δ holoenzymes resumes 3 nt downstream of the lesion (**Figure 5C**). Utilization of DDT for extension may be promoted in the absence of intrinsic proofreading by pol δ as inactivation of the 3′→5′ exonuclease activity of human pol δ significantly decreases the extension efficiency (by 28.2 ± 2.93 %, **Figure 6A**) leading to a significant increase (17.8 ± 2.44%, **Figure 6B**) in pol δ dissociation during this dNTP in corporation step.

### O6MeG in lagging strand templates

Similar to 8oxoG and Tg, human pol δ holoenzymes are also remarkably efficient at inserting a dNTP opposite O6MeG (insertion efficiency = 79.6 ± 0.617 %, **Figure 11A**) such that 78.2 ± 0.605 % of progressing pol δ holoenzymes that encounter an O6MeG complete insertion prior to pol δ dissociating (**Figure 11B**). Furthermore, like 8oxoG and Tg, intrinsic proofreading visibly contributes (12.6 ± 1.04 %) to this unexpectedly high insertion efficiency (**Figure 8A**). Previous studies indicated that, in the presence of PCNA and RPA, human pol δ primarily inserts either the correct dCTP or the incorrect dTTP opposite 8oxoG, and these events occur with equal probability (14). In the present study, this equates to ∼39.2% of all encounters between progressing pol δ holoenzymes and 8oxoG lesions resulting in O6MeG:T mismatches and ∼39.2% yielding “correct” O6MeG:C base pairs. Altogether, this suggests that the overwhelming majority of O6MeG lesions in lagging strand templates are *initially* replicated by pol δ, rather than a DDT pathway, creating a heterogenous population of nascent DNA that may elicit a variety of downstream responses during DNA replication. For O6MeG:C base pairs, pol δ faithfully completes insertion and DDT would only be utilized, if at all, to complete extension as dissociation of pol δ is most prevalent at this dNTP incorporation step (**Figure 11B**) and unperturbed progression of pol δ holoenzymes resumes 2 nt downstream of the lesion (**Figure 7C**). Utilization of DDT for extension may be promoted in the absence of intrinsic proofreading by pol δ as inactivation of the 3′→5′ exonuclease activity of human pol δ significantly decreases the extension efficiency (by 33.1 ± 1.02 %, **Figure 8A**) leading to a significant increase (19.9 ± 0.294%, **Figure 8B**) in pol δ dissociation during this dNTP in corporation step. For O6MeG:T mismatches, the mismatched dTMP opposite the O6MeG lesion must be excised and the O6MeG accurately “re-replicated” to avoid fixed G:C → A:T transversion mutations. This may occur via multiple pathways that depend on the extent of pol δ holoenzyme progression downstream (3′) of an O6MeG lesion and/or the activation of DDT pathways. We discuss two possibilities below.

O6MeG:T mismatches within dsDNA are excellent substrates for mismatch repair (MMR) pathway (66,67). MMR is physically and functionally coupled to DNA replication, excises nascent DNA containing a mismatched dNMP, and then re-replicates the re-exposed template ssDNA sequence. MMR requires PCNA, which is left behind at P/T junctions aborted by pol δ, and significant nascent DNA downstream (3′) of the mismatch (68,69). In the present study, over 60% (63.1 ± 0.819 %) of progressing pol δ holoenzymes that insert a dNTP opposite O6MeG subsequently extend the nascent DNA at least 19 nt downstream prior to pol δ dissociation. This suggests that the majority of the O6MeG:T mismatches generated by progressing pol δ holoenzymes are subsequently converted by the holoenzymes into substrates for MMR. The remainder may be extended to sufficient lengths for MMR by pol δ (in subsequent binding encounters) or DDT pathways, if necessary. However, template ssDNA sequences exposed during MMR are primarily replicated by human pol δ holoenzymes, which are remarkably efficient (**Figure 11A**) and error-prone at replicating O6MeG lesions (mutation frequency = 50%) (14,70). Hence, O6-methylated guanines must be repaired prior to their re-replication by pol δ holoenzymes to ensure that O6MeG:T mismatches are not re-formed. Failure to do so can lead to another round of MMR and potentially enter the afflicted genomic DNA sequence into futile MMR cycles that ultimately result in dsDNA breaks (71). O^6^-methylguanine DNA methyltransferase (MGMT), also known as O^6^-alkylguanine-DNA alkyltransferase (AGT), stoichiometrically removes alkyl adducts at the O6-position of guanine by direct reversal. Interestingly, human MGMT is equally efficient in reversing O6 methylation of guanine nucleobases residing in dsDNA and ssDNA (72-78). Thus, O6-methylation reversal may occur before or after MMR-dependent excision of the nascent DNA containing the mismatched dTMP. Upon restoration of the native guanine, the exposed template ssDNA sequence can be faithfully replicated by a pol δ holoenzyme.

Alternatively, P/T junctions aborted by pol δ shortly downstream of an O6MeG:T mismatch may be re-engaged by a factor with 3′ → 5′ exonuclease activity that degrades the nascent DNA to ultimately excise a mismatched dTMP opposite O6MeG. PCNA and RPA, both of which are left behind at P/T junctions aborted by pol δ, may recruit such factors. In the present study, the 3′ → 5′ exonuclease activity of human pol δ significantly promotes bypass of O6MeG lesions (**Figure 8A, Table 3**) by intrinsically proofreading 1 or both steps of lesion bypass (i.e., insertion and extension). Thus, pol δ may excise a mismatched dTMP opposite O6MeG during subsequent binding encounters. Excision may also be carried out by WRN or a WRN•Pol δ complex (described above). In support of this possibility, siRNA-mediated knockdown of WRN in human cells increases the sensitivity to O6MeG and the frequency of G:C → A:T transversion mutations induced by O6MeG (79,80). In this MMR-independent scenario, any PCNA-dependent DNA polymerase may re-replicate the offending O6MeG lesion and all human DNA polymerases analyzed to date (pols β, δ, η, ι, κ, and ν) replicate O6MeG with remarkably low fidelity. This suggests that O6 methylated guanines must still be repaired by MGMT prior to their re-replication to avoid regeneration and subsequent re-processing of O6MeG:T mismatches (14,81-87). Pol δ is likely selected for replication of the resultant native DNA sequence due to its unique interaction with RPA (88) and its superior affinity for PCNA encircling a P/T junction compared to other DNA polymerases (28,89).

### εA in lagging strand templates

In contrast to 8oxoG, Tg, and O6MeG, human pol δ holoenzymes are very inefficient at inserting a dNTP opposite εA (insertion efficiency = 3.11 ± 0.568, **Figure 11A**) such that only 3.06 ± 0.558 % of progressing pol δ holoenzymes that encounter an εA complete insertion prior to pol δ dissociating (**Figure 11B**). Furthermore, human pol δ holoenzymes are incapable of extending from a dNMP inserted opposite εA. Finally, intrinsic proofreading by pol δ does not contribute to the aforementioned behaviors (**Figure 10**). Altogether, this suggests that εA are very strong blocks to pol δ holoenzyme progression and, consequently, nearly all εA lesions in lagging strand templates are replicated by a DDT pathway. This agrees with a previous *ex vivo* study (90). It is also possible that upon dissociation of pol δ during insertion opposite εA or extension from an εA base pair, the offending lesion is subsequently repaired via direct reversal and the restored native adenine is then replicated by a pol δ holoenzyme (91-93). This is currently under investigation.

## Data Availability

## Funding

## Acknowledgements

We would like to thank all members of the Hedglin lab for their efforts in reviewing/proofreading the current manuscript.

## Conflict of Interest Disclosure

The authors declare that they have no conflicts of interest with the contents of this article^1^

## AUTHOR CONTRIBUTIONS

K.G.P. and R.L.D. expressed, purified, and characterized all proteins. R.L.D. and J.A.C. performed the experiments and analyzed the data. M.H. designed the experiments and analyzed the data. M.H. wrote the paper.

## Notes

+ Supported by a Benkovic Award for Undergraduate Research

### Competing Interest Statement

The authors have declared no competing interest.

