## Supporting Information for "High resolution studies of DNA lesion bypass by human DNA polymerase δ holoenzymes"

[+]Supported by a Benkovic Award for Undergraduate Research

**Affiliations**:
[1]Department of Chemistry, The Pennsylvania State University, University Park, PA 16802

### Experimental Methods

*FRET-based assay to monitor extent of RFC-catalyzed loading of PCNA.* All experiments were performed at room temperature (23 ± 2 °C) in 1X Replication Buffer (25 mm HEPES, pH 7.5, 10 mm $Mg(OAc)_2$, 125 mm KOAc) supplemented with 1 mm DTT, and the final ionic strength was adjusted to physiological (200 mm) by the addition of appropriate amounts of KOAc. All measurements were done in 16.100F-Q-10/Z15 sub-micro fluorometer cells (Starna Cells) in a Horiba Scientific Duetta-Bio fluorescence/absorbance spectrometer. Excitation and emission slit widths are each set to 5 nm, unless indicated otherwise. Reaction solutions are excited at 514 nm and the fluorescence emission intensities ($I$) are simultaneously monitored at 563 nm ($I_{563}$, Cy3 FRET donor fluorescence emission maximum) and 665 nm ($I_{665}$, Cy5 FRET acceptor fluorescence emission maximum) over time, recording $I$ every 0.17 s. For each time point, $E_{FRET}$ is calculated where $E_{FRET} = \frac{I_{665}}{I_{665}+I_{563}}$. $E_{FRET}$ values for complete loading of PCNA onto a P/T junction were calculated based on a published FRET-based assay(1-9). A solution containing 110 nM of a Bio-Cy3P/T DNA (**Figure S1**), 440 nM neutravidin and 1 mM ATP is pre-incubated with RPA (330 nM heterotrimer). Then, Cy5-PCNA (100 nM homotrimer) is added. Finally, RFC (100 nM heteropentamer) is added, the resultant solution is mixed via pipetting, and $E_{FRET}$ is monitored beginning 10 s after the addition of RFC. Data is plotted as a function of time after RFC addition with time courses adjusted for the time between the addition of RFC and the recording of $E_{FRET}$ ($\Delta t$ = 10 s).

### Supporting Figures

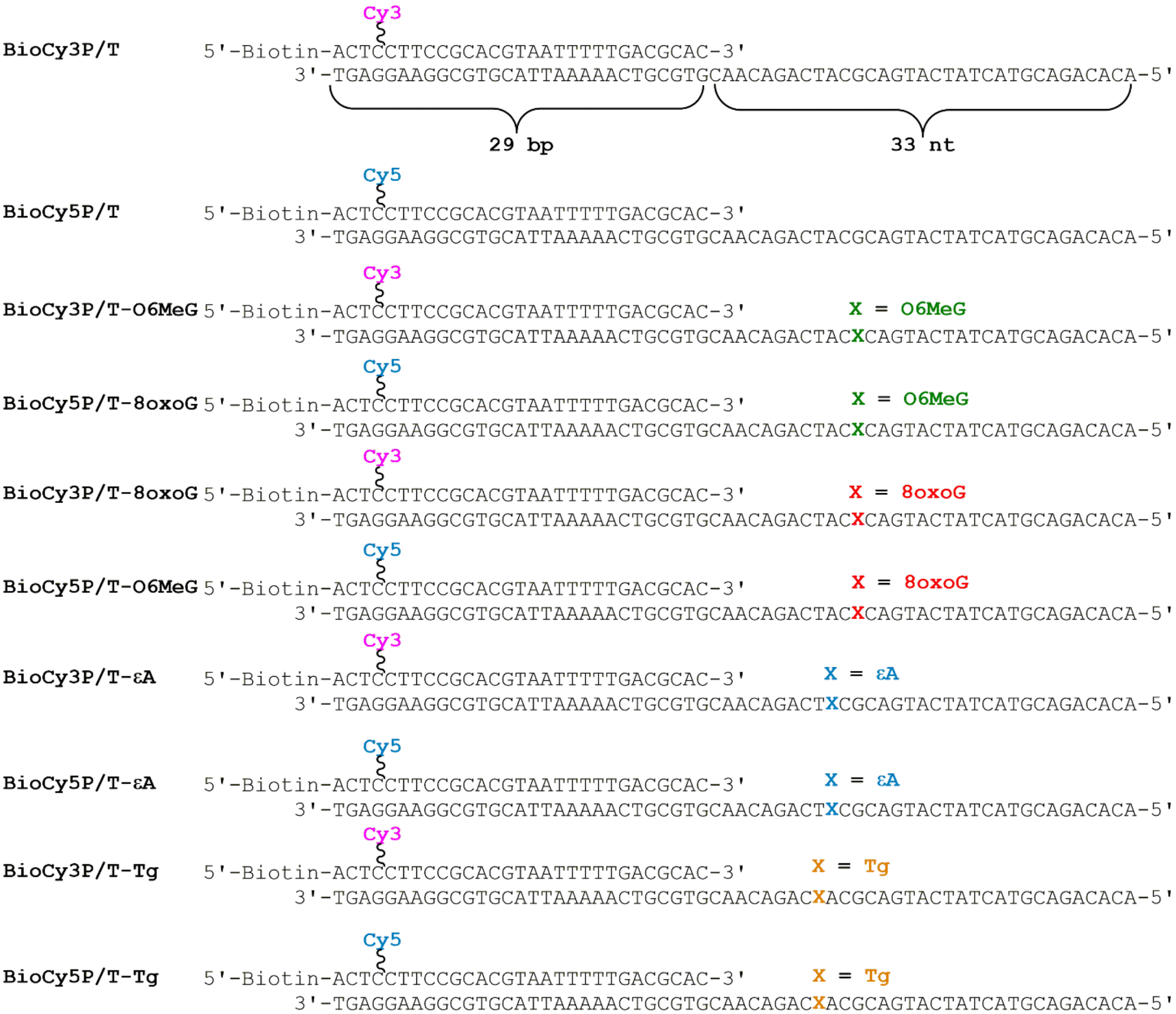


**Figure S1**. DNA substrates utilized in this study. For all P/T DNA substrates, the sequences and lengths (29 bp) of the double strand DNA (dsDNA) regions are identical and in agreement with the requirements for assembly of a single PCNA ring onto DNA by RFC (1,4,7). The ssDNA regions adjacent to the 3'-end of the P/T junctions accommodate 1 RPA molecule (10-12). RPA prevents loaded PCNA from sliding off the ssDNA end of the substrate (4). When pre-bound to neutravidin, the biotin attached to the 5' end of a primer strand prevents loaded PCNA from sliding off the dsDNA end of the substrate.

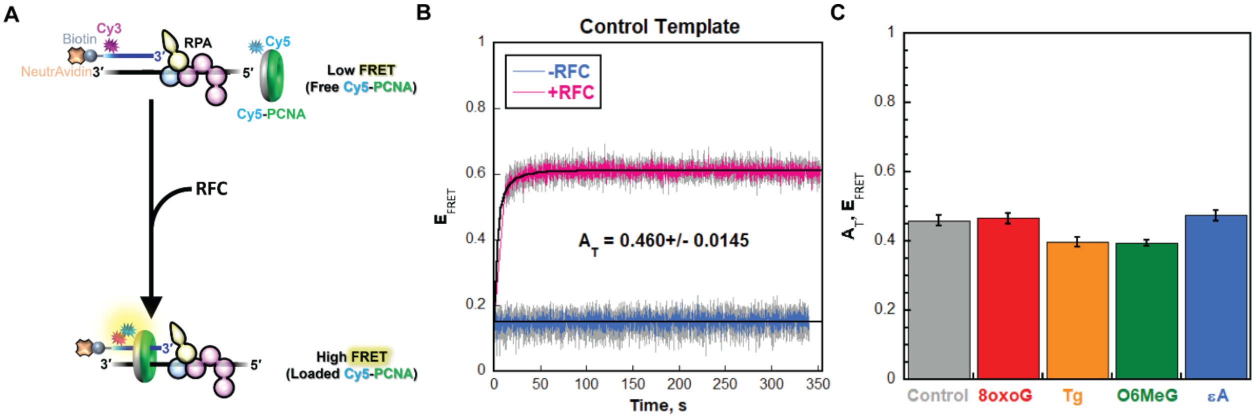


**Figure S2.** RFC-catalyzed loading of PCNA onto DNA. (**A**) Schematic representation of the assay. The DNA substrates are P/T DNA substrates (**Figure S1**) in which the primer contains an internal Cy3 (FRET donor) 4 nt from its 5′ terminus. The "back face" of PCNA (shown in grey) is labeled with a Cy5 (FRET acceptor). Cy5-PCNA is loaded onto the DNA substrate by the human clamp loader, RFC, such that the Cy5 FRET donor on the "back face" of PCNA faces the Cy3 FRET donor near the 5′ terminus of the primer strand and the "front face" of PCNA (shown in green) is oriented towards the P/T junction where DNA synthesis emanates from. Cy5-PCNA is loaded by RFC in the presence of excess RPA and $E_{FRET}$ is monitored over time. Under these conditions, all Cy5-PCNA is loaded onto a native (i.e., undamaged) BioCy3P/T DNA substrate (**Figure S1**) by RFC and stabilized by RPA and the biotin/neutravidin blocks that prevent diffusion of PCNA off of the DNA (1-9). (**B**) FRET data observed in the absence and presence of RFC for the BioCy3P/T-R DNA substrate ("control"). $E_{FRET}$ is plotted as function of time after RFC is added. Each $E_{FRET}$ trace is the average of at least three independent traces and the standard error is shown in grey. FRET is not observed when RFC is omitted and the signal remains constant. For these conditions, the traces are fit to a flat line. $E_{FRET}$ increases with time only when RFC is included. For these conditions, the traces are fit to a double exponential rise where the sum of the amplitudes for the two phases ($A_T$, shown) indicates the change in $E_{FRET}$ observed when Cy5-PCNA is stably loaded onto the BioCy3P/T-R DNA substrate. Thus, $A_T$ directly reports on the extent of stable assembly of PCNA onto DNA. (**C**) $A_T$ data for the BioCy3P/T, BioCy3P/T-8oxoG, BioCy3P/T-Tg, BioCy3P/T-O6MeG, and BioCy3P/T-εA DNA substrates are shown in grey, red, orange, green, and blue, respectively. The $A_T$ value observed for the control DNA substrate (i.e., native/undamaged) agrees with that observed previously (1-9), indicating that all Cy5-PCNA is loaded onto the P/T junction and stabilized. The $A_T$ values observed for the DNA substrates containing a DNA lesion at least 9 nt downstream of the P/T junction are identical to that observed for control DNA substrate. This indicates that DNA lesions do not affect the stable assembly of PCNA onto a P/T junction upstream of the lesion.

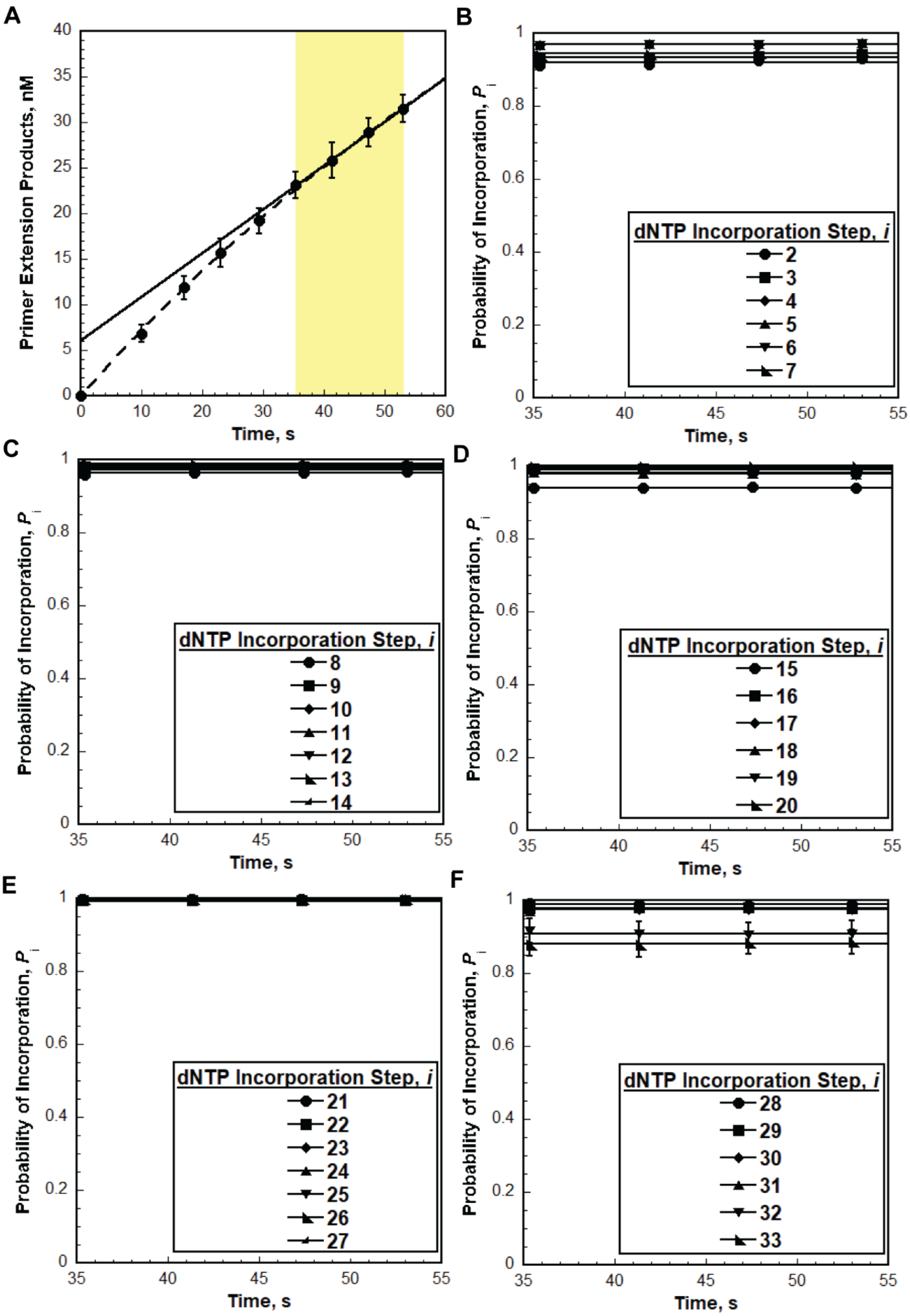
A
Primer Extension Products, nM
Time, s
B
Probability of Incorporation, $P_i$
dNTP Incorporation Step, i
2
3
4
5
6
7
Time, s
C
Probability of Incorporation, $P_i$
dNTP Incorporation Step, i
8
9
10
11
12
13
14
Time, s
D
Probability of Incorporation, $P_i$
dNTP Incorporation Step, i
15
16
17
18
19
20
Time, s
E
Probability of Incorporation, $P_i$
dNTP Incorporation Step, i
21
22
23
24
25
26
27
Time, s
F
Probability of Incorporation, $P_i$
dNTP Incorporation Step, i
28
29
30
31
32
33
Time, s

**Figure S3**. Monitoring primer extension by pol δ holoenzymes during a single binding encounter with a P/T DNA substrate. (**A**) Quantification of the (total) primer extension products for pol δ holoenzymes on the native (i.e., undamaged) P/T DNA substrate (BioCy3P/T, **Figure S1).** Data is identical to that displayed in **Figure 2C** in the main text. Each data point represents the average ± S.E.M. of 3 independent experiments. Data is plotted as a function of time (after the addition of pol δ) and displays "burst" kinetics. Data points within the "linear" phase (highlighted in yellow) are fit to a liner regression where the Y-intercept (in nM) represents the amplitude of the "burst" phase and the slope represents the initial velocity (in nM/min) of the "linear" phase. (**B - F**) Processivity of pol δ holoenzymes at single nucleotide resolution. The probability of incorporation ($P_i$) values for each dNTP incorporation step ($i$) beyond the first incorporation step ($i = 2$) are calculated for each time point within the "linear phase" of the reaction (indicated in panel **A**) for the experiments depicted in **Figure 2** and plotted as a function of time. The results for $i = 2$ to $i = 7$, $i = 8$ to $i = 14$, $i = 15$ to $i = 20$, $i = 21$ to $i = 37$, and $i = 28$ to $i = 33$ are depicted in panels **B**, **C**, **D**, **E**, and **F**, respectively, and each data point represents the average ± S.E.M. of 3 independent experiments. For each dNTP incorporation step ($i$), $P_i$ values remain constant within this incubation time. Identical behavior was observed for all P/T DNA substrates utilized in this study with all forms of pol δ.

| | | Pol δ | | Pol $\delta^{Exo-}$ | |
|---|---|---|---|---|---|
| | | Avg | StdErr | Avg | StdErr |
| Insertion Probability | G | 0.983 | 3.65E-04 | 0.964 | 1.75E-03 |
| | 8oxoG | 0.842 | 6.34E-03 | 0.731 | 5.98E-03 |
| Insertion Efficiency (%) | | 85.7 | 0.646 | 75.8 | 0.635 |
| Extension Probability | G | 0.988 | 5.18E-04 | 0.967 | 1.97E-03 |
| | 8oxoG | 0.432 | 0.0144 | 0.364 | 0.0143 |
| Extension Efficiency (%) | | 43.7 | 1.45 | 37.7 | 1.48 |
| Bypass Probability | G | 0.971 | 6.06E-04 | 0.932 | 3.56E-03 |
| | 8oxoG | 0.364 | 0.0148 | 0.266 | 9.55E-03 |
| Bypass Efficiency (%) | | 37.5 | 1.52 | 28.6 | 1.030 |

**Table S1**. Efficiency of human pol δ holoenzymes replicating 8oxoG

| | | WT | | Exo- | |
|---|---|---|---|---|---|
| | | Avg | StdErr | Avg | StdErr |
| Insertion Probability | T | 0.979 | 9.73E-04 | 0.969 | 2.05E-03 |
| | Tg | 0.775 | 6.56E-03 | 0.731 | 1.09E-02 |
| Insertion Efficiency (%) | | 79.1 | 0.674 | 75.4 | 1.133 |
| Extension Probability | T | 0.983 | 8.48E-04 | 0.974 | 1.96E-03 |
| | Tg | 0.408 | 2.65E-02 | 0.129 | 1.12E-02 |
| Extension Efficiency (%) | | 41.5 | 2.696 | 13.3 | 1.150 |
| Bypass Probability | T | 0.963 | 1.78E-03 | 0.944 | 3.89E-03 |
| | Tg | 0.316 | 0.0229 | 0.094 | 0.0068 |
| Bypass Efficiency (%) | | 32.9 | 2.38 | 10.0 | 0.72 |

**Table S2**: Efficiency of human pol δ holoenzymes replicating Tg

| | | WT | | Exo- | |
|---|---|---|---|---|---|
| | | Avg | StdErr | Avg | StdErr |
| Insertion Probability | G | 0.983 | 3.65E-04 | 0.964 | 1.75E-03 |
| | O6MeG | 0.782 | 6.05E-03 | 0.646 | 7.97E-03 |
| Insertion Efficiency (%) | | 79.6 | 0.617 | 67.0 | 0.835 |
| Extension Probability | G | 0.988 | 5.18E-04 | 0.967 | 1.97E-03 |
| | O6MeG | 0.854 | 2.44E-03 | 0.516 | 9.47E-03 |
| Extension Efficiency (%) | | 86.5 | 0.251 | 53.3 | 0.986 |
| Bypass Probability | G | 0.971 | 6.06E-04 | 0.932 | 3.56E-03 |
| | O6MeG | 0.668 | 0.0060 | 0.334 | 0.0102 |
| Bypass Efficiency (%) | | 68.8 | 0.62 | 35.8 | 1.10 |

**Table S3**: Efficiency of human pol δ holoenzymes replicating O6MeG

| | | WT | | Exo- | |
|---|---|---|---|---|---|
| | | Avg | StdErr | Avg | StdErr |
| Insertion Probability | A | 0.983 | 8.48E-04 | 0.974 | 1.96E-03 |
| | εA | 0.031 | 5.58E-03 | 0.042 | 3.01E-03 |
| Insertion Efficiency (%) | | 3.11 | 0.568 | 4.34 | 0.309 |
| Extension Probability | A | 0.972 | 1.15E-03 | 0.951 | 2.28E-03 |
| | εA | 0.000 | 0.00E+00 | 0.000 | 0.00E+00 |
| Extension Efficiency (%) | | 0.0 | | 0.0 | |
| Bypass Probability | A | 0.955 | 1.91E-03 | 0.926 | 4.04E-03 |
| | εA | 0.000 | 0.0000 | 0.000 | 0.0000 |
| Bypass Efficiency (%) | | 0.0 | | 0.0 | |

**Table S4**: Efficiency of human pol δ holoenzymes replicating εA.
